# Evolution of crop phenotypic spaces through domestication

**DOI:** 10.1101/2025.10.11.681794

**Authors:** Arthur Wojcik, Harry Belcram, Agnès Rousselet, Manon Bouët, Marie Brault, Pierre Serin, Augustin Desprez, Renaud Rincent, Andreas Peil, Karine Henry, Cécile Marchal, Thierry Lacombe, Sylvain Glémin, Karine Alix, Pierre R. Gérard, Domenica Manicacci, Yves Vigouroux, Catherine Dogimont, Maud I. Tenaillon

**Affiliations:** Université Paris-Saclay, INRAE, CNRS, AgroParisTech, GQE - Le Moulon, EMR 9005, 91190, Gif-sur-Yvette, France; INRAE Avignon, GAFL - Génétique et Amélioration des Fruits et Légumes, 84143, Montfavet, France; Université de Montpellier, IRD, CIRAD, DIADE, 34394, Montpellier, France; Université Paris-Saclay, INRAE, AgroParisTech, GQE - Le Moulon, 91190, Gif-sur-Yvette, France; Julius Kühn-Institut, 01326, Pillnitz, Germany; Florimond-Desprez, Cappelle-en-Pévèle, 59242, France; Grapevine Biological Resources Center, INRAE, Université de Montpellier, Unité Expérimentale Domaine de Vassal, 34340, Marseillan, France; Université Montpellier, CIRAD, INRAE, Institut Agro, AGAP Institut, 34398, Montpellier, France; Université de Rennes, CNRS, ECOBIO, 35000, Rennes, France; Department of Ecology and Genetics, Evolutionary Biology Centre, Uppsala University, 75236, Uppsala, Sweden

**Keywords:** Domestication index, Divergent selection, Phenotypic convergence, Life-history traits, Near infrared spectra, Trait correlations

## Abstract

- We used domestication as an *in vivo* replicated experiment to investigate how divergent selection has shaped the evolution of multivariate phenotypic spaces.
- We measured 11 to 57 qualitative and quantitative traits in 13 species, either unique or shared between species, and established a framework for cross-species comparisons.
- Our results revealed significant convergence that translated into a cross-species domestication syndrome. Most species exhibited a reduction of the multivariate phenotypic space during domestication. We brought evidence that Near Infrared spectra measured on leaves reflect phenotypic evolution unrelated with domestication, enabling its use as a control for sampling effects across species. Building on this, we developed a multivariate Phenotypic Divergence Index (mPDI) to rank species by the extent of phenotypic divergence under domestication. We found a high disjunction of wild and domestic phenotypic spaces in all species. Neither the mPDI nor the relative size of wild versus domestic multivariate phenotypic spaces was influenced by the domestication timing or mating system. Lastly, we observed a progressive decoupling of trait correlations with increasing time since domestication.
- In addition to introducing a new index that can be applied for cross-species comparisons, our study uncovers recurring patterns shared among species, pointing to general principles underlying plant domestication.

## Introduction

Domestication is a process through which organisms evolve in response to human-altered habitats (Lord et al., 2025). One of the outcomes of this process is phenotypic divergence between the forms, resulting from disruptive selection between wild populations shaped by natural selection in their native habitats and domestic populations shaped by the combined effects of natural selection and adaptation to anthropogenic niches (Glémin & Bataillon, 2009; Ross-Ibarra et al., 2007), which are characterized by recurrent disturbance, habitat simplification, redistribution of resources, and relaxed biotic interactions such as competition and predation (Ellis 2015). From an evolutionary standpoint, plant domestication is a recent process that began during the neolithic revolution (Weisdorf, 2005) with the earliest known domestications occurring ∼12,000 years ago in the Fertile Crescent (Barker, 2006). Although debates persist about the intentionality of early selection (Abbo & Gopher, 2020), archaeological evidence from the few crop species for which extensive temporal series are available supports a protracted model of domestication, with the slow fixation of domestic traits over thousands of generations as shown in einkorn wheat, rice and barley (Purugganan & Fuller, 2011). After domestication, crop expanded through trade and seed exchange, leading to the emergence of a genetic structuring into groups, well described in e.g., maize or pearl millet (Burgarella et al., 2018; Tenaillon & Charcosset, 2011).

Over 2500 plants species belonging to 173 botanical families have undergone cultivation or domestication — providing an upper bound for the number of domesticated plants (reviewed in Dirzo & Raven, 2003). Human selection has targeted a diversity of organs (e.g., flower, fruit, seed, tuber) and usages (e.g., food, fiber, ornamentals, medicine). Phenotypic changes that often arise through parallel evolution across species under domestication are collectively referred to as the phenotypic domestication syndrome. These changes include traits that facilitate cultivation, harvesting, and consumption and differ between crop-types e.g., between herbs and woody plants or between vegetatively and sexually propagated crops, for example (Fuller et al., 2023). Domestication traits are well-documented in annual crops. In cereals, for example, they include reduced tillering, loss of seed shattering, and increased ear and seed size (Glémin & Bataillon, 2009; Tenaillon & Manicacci, 2011). While wild progenitors of crops tend to be fast-growing, with high resource-capture ability (de Casas et al., 2025; Milla et al., 2015), domestication has favored greater above-ground biomass, enhancing competitiveness in cultivated environments, though consistent effects on leaf physiological traits have not been demonstrated (reviewed in Milla, 2023). Finally, domestication has occasionally modified life-history traits, such as an increase in selfing rate in sunflower, cacao or grapevine (Dempewolf et al., 2012). Compared to annual species, perennial crops are characterized by extended juvenile phases, fewer sexual generations, and a reliance on clonal propagation, which together make them younger domesticates (Gaut et al., 2015). As in annual seed crops, domestication has favored a more compact and simplified plant architecture with reduced branching, increased edible organs, loss of bitter compounds and reduced sexual reproduction for vegetatively reproduced crop (Denham et al., 2020; Miller & Gross, 2011). Hence, convergence of domestication traits across species within-crop types, well documented in cereals (Tenaillon & Manicacci, 2011), seems to extend to more distant taxa (Meyer et al., 2012) but has yet to be tested.

The phenotypic domestication syndrome is accompanied by a molecular domestication syndrome (Burban et al., 2022), primarily characterized by a loss of genetic diversity due to bottlenecks and linked selection at key domestication genes (Meyer & Purugganan et al. 2013; Gaut et al., 2015). These combined processes have likely resulted in a loss of additive genetic variance underlying quantitative traits targeted by selection, as shown in maize (Yang et al., 2019). Thus, one might expect a reduction of the domestic phenotypic space—that is, a reduction in the range of observable trait variation expressed by domesticated forms—as divergence from wild populations increases. However, crop expansion into new environments and selection for local preferences (Bradshaw, 2016; Meyer & Purugganan, 2013) generated a range of new phenotypes, not directly linked to domestication, which would partially offset this reduction. To test whether domestication itself has reduced phenotypic space, it is therefore essential to apply a careful sampling that minimizes the inflating effects of post-domestication processes such as genetic structuring and modern breeding.

Another intriguing question arises from the concerted evolution of domestication traits whose determinism involves complex gene regulatory networks (Studer et al., 2017). Several studies indicate that domestication has led to a deep rewiring of these networks as shown by the reorganization of co-expression networks (e.g., in cotton Rapp et al., 2010), common bean (Bellucci et al., 2014), and tomato (Sauvage et al., 2017)). Simulations suggest that the combined effects of demography and selection during domestication result in a redistribution of genetic correlations, leading to less distinct clusters and a moderate increase of these correlations (Burban et al., 2022). Studies on maize and teosintes indeed showed an overall change in the structure of the correlation matrix, but the findings were less clear for reproductive traits directly affected by domestication, with some analyses indicating conservation of correlations in the two forms. Contrary to the model predictions, teosintes generally exhibit more significant and stronger correlations than maize (Yang et al., 2019). Model predictions therefore need to be further tested across a wider range of species.

In sum, although individual domestication traits have now been described in many systems, our understanding of the evolution of phenotypic spaces through the domestication process, and the underlying key evolutionary mechanisms, remains superficial. This is partly due to a lack of unifying statistical framework able to integrate data from different species, where various and often distinct traits are experimentally measured, including traits of different types (quantitative and qualitative) and scales. The approach we propose here is to build on multivariate analysis to transform measured traits into statistically comparable objects across species; and to use a control trait, a priori unrelated with domestication and comparable across species, to account for sampling heterogeneity across species providing a baseline of wild and domestic phenotypic divergence largely independent of domestication. Using Near Infrared Spectra (NIRS) measured on leaves as the control trait, we derived a multivariate Phenotypic Divergence Index (mPDI), which quantifies the degree of phenotypic divergence between wild and domesticated forms in each species.

By applying this unifying framework to 13 wild/domesticated pairs (13 species) including a range of domestication dates from ancient to more recent, different mating systems - outcrossers and selfers, and various life cycles from annual to perennial, we addressed the following questions: (1) Is there a significant phenotypic convergence across species? (2) How did phenotypic spaces evolve during domestication? (3) Do domestication dates and mating systems influence the mPDI? (4) Are there consistent patterns in trait correlations throughout the domestication process?

## Material and Methods

### Plant Material

Our study examined 13 domestic plant species and the wild progenitors from which they were derived through domestication: African rice (*Oryza glaberrima* Steud.), apple (*Malus domestica* Borkh.), cabbage (*Brassica oleracea* L.), common bean (*Phaseolus vulgaris* L.), eggplant (*Solanum melongena* L.), einkorn wheat (*Triticum monoccocum* L.), foxtail millet (*Setaria italica* (L.) P. Beauv.), grapevine (*Vitis vinifera* L.), maize (*Zea mays*), melon (*Cucumis melo* L.), pearl millet (*Pennisetum glaucum* (L.) R. Br.), sugar beet (*Beta vulgaris* L. subsp. *vulgaris*) and tomato (*Solanum lycopersicum* L.).

These 13 wild–domestic pairs were selected because they are diploid and together encompass a broad domestication timeline, ranging from the earliest domestication of einkorn wheat approximately 9,500 years before present (YBP) to the more recent domestication of eggplant around 2,000 YBP (Table S1), with times also expressed in generations (Fig. 1a). The pairs also span phylogenetically distant taxa across eight botanical families (Fig. 2a; Table S1). Importantly, the 13 species exhibit a diversity of life-cycles from annual, biennial (e.g., cabbage) to perennial (e.g., grapevine), with sugar beet transitioning from annual to biennial during domestication (Table S1, Fig. 1a). And, finally, the pairs display a range of reproductive systems from highly self-fertilizing species (selfers) such as foxtail millet to obligate outcrossers such as apple tree (Table S1). Most species have hermaphroditic flowers, except for maize which is monoecious, grapevine that transitioned from dioecy to monoecy during domestication, and melon from monoecy to andromonoecy (Table S1).

**Figure 1:**
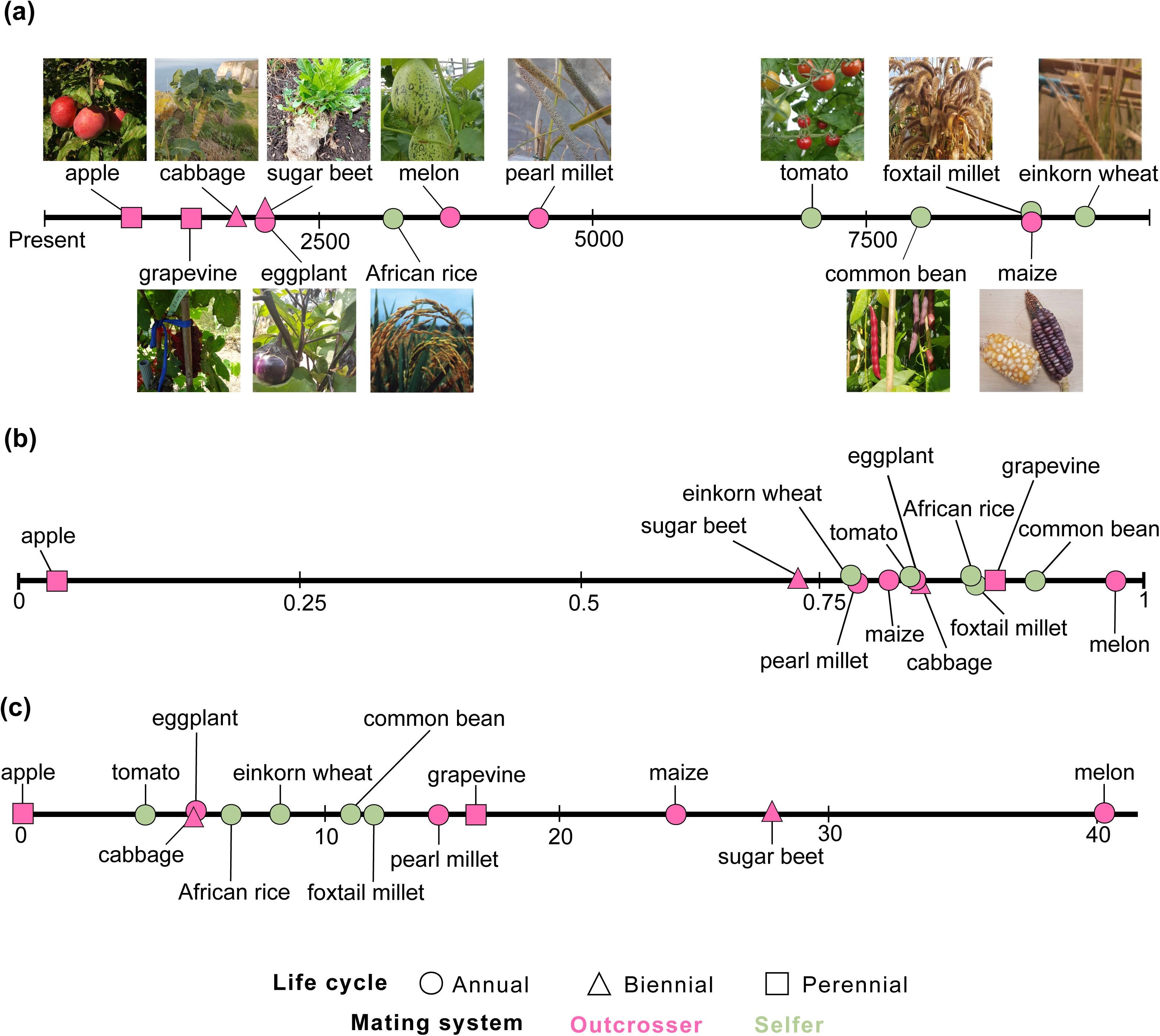
Domestication timing and multivariate phenotypic divergence index (mPDI) for the 13 species. Domestication timing (Table S1) is indicated in generations (a). Species are ranked by their mPDI (b) and the mPDI/*Pillai_control_* (c).

**Figure 2:**
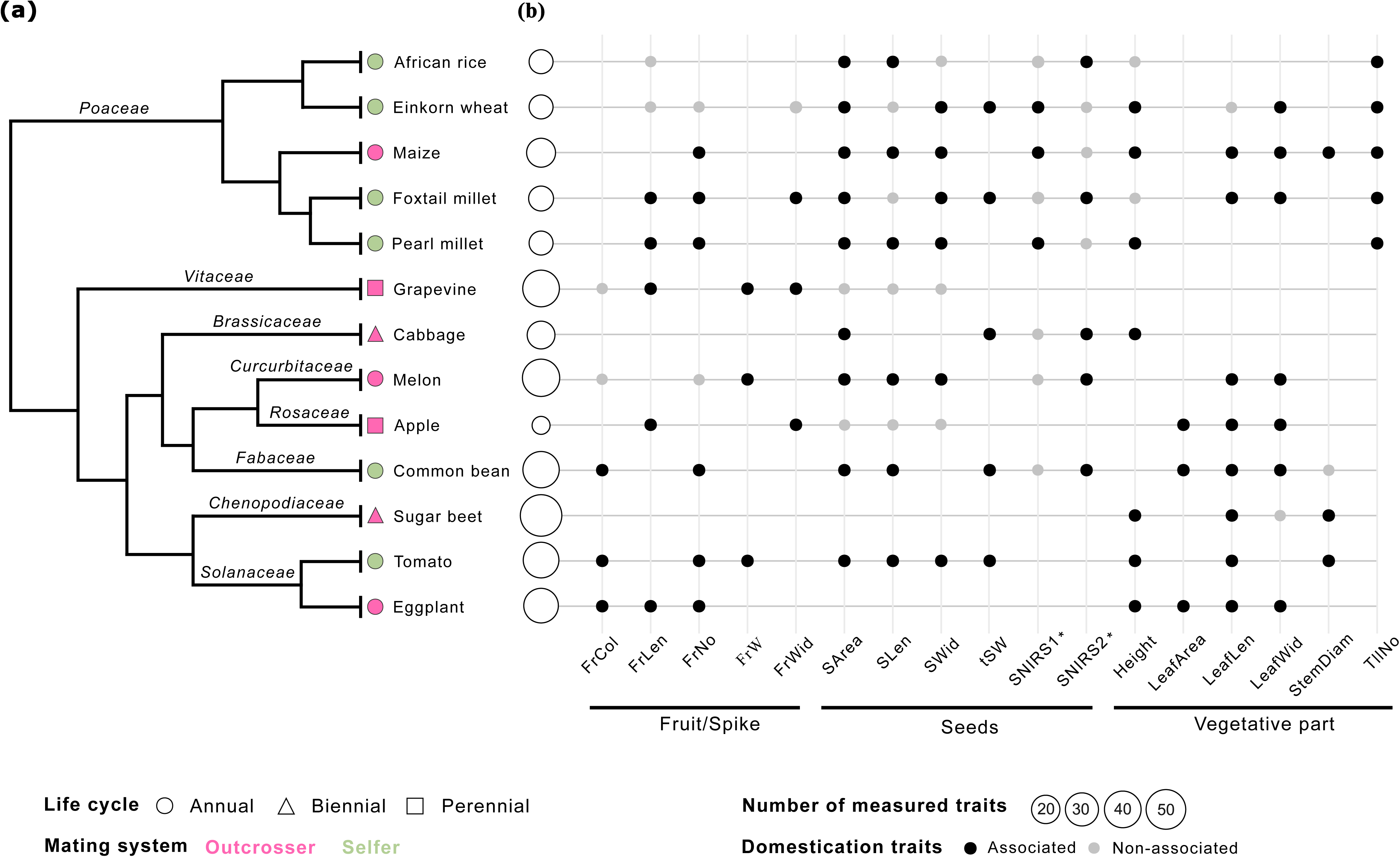
Domestication-associated traits shared by multiple species. Phylogenetic relationships of the 13 species (adapted from Milla et al., 2020) (a) and shared domestication-associated traits in at least three species (b). The length of the tree branches is not to scale. *: SNIRS1 and SNIRS2 cannot be considered homologous traits among species because NIRS relies on the first two axes of a PCoA, with potentially different wavelengths contributing unequally to each axis across species. FrCol: Color of fruit; FrLen: Fruit length; FrNo: Number of fruits per plant; FrW: Fruit weight; FrWid: Fruit Width; SArea: Seed area; SLen: Seed length; SWid: Seed width; tSW: Thousand seed weight; SNIRS1: First axis of PCoA on NIRS_seed_; SNIRS2: Second axis of PCoA on NIRS_seed_; Heigth: Height of plant; LeafArea: Leaf area; LeafLen: Leaf length; LeafWid: Leaf width; StemDiam: Stem Diameter; TllNo: Tiller number per plant.

To facilitate a comparative analysis across species, we aimed to apply consistent sampling criteria within each species. For species where two domestication centers were recorded, we focused on a single domestication center and the corresponding wild gene pool: South America for tomato, Mesoamerica for common bean, and Sudan for melon. For beet, we focused on sugar beet whose domestication happened later than for leaf beet (Table S1). In each species, we chose 17 to 20 wild populations (wild accessions) to represent the wild gene pool and 16 to 22 landraces (domestic accessions) to represent the domestic gene pool (Table S2) except for melon and eggplant for which the wild sampling encompassed 8 and 13 populations, respectively. For the domestic gene pool, we focused on traditional landraces in order to capture the domestication process itself while minimizing the effects of more recent breeding.

In addition, we minimized genetic structuring among individuals to avoid confounding effects of genetic group in phenotypic analyses. To this end, we relied on established domestication centers (Table S1) and previous genetic structuring analyses for sampling both wild and domestic accessions in nine of the systems: African rice (Cubry et al., 2018), cabbage (Maggioni et al., 2020), common bean (Bellucci et al., 2023), eggplant (Ranil et al., 2017), einkorn wheat (Adhikari et al., 2022), foxtail millet (Le Thierry d’Ennequin et al., 2000), maize (Aguirre-Liguori et al., 2017; van Heerwaarden et al., 2011), melon (Zhao et al., 2019), pearl millet (Burgarella et al., 2018), sugar beet (Mangin et al., 2015) and tomato (Razifard et al., 2020). Perennials were exceptions in this sampling design, as we relied on existing and accessible tree collections. For grapevine, whose domestication center is in South Caucasus, we focused on the western Mediterranean region, where a greater number of wild and domestic accessions were available (Laucou et al., 2018). For apple, the domestic sample was confined to European traditional cultivars, while wild samples originate from Central Asia (Cornille et al., 2012). For cabbage, where the exact European domestication center is still debated (Mabry et al., 2021), we focused on wild and domestic accessions from western Europe (France and United Kingdom).

### Plant phenotyping

Except for seed germination rate which required to evaluate many seeds per accession, plant phenotyping was conducted using a three randomized-replicate design for all but four species (melon, cabbage, apple, grapevine), with each replicate containing a single individual from each wild accessions and each landrace for a total of maximum 120 individuals – 3x40 initial individuals (Table S3). For melon and cabbage, we measured a single individual per accession in a randomized design (i.e., no replicate). For apples, we also evaluated a single individual per accession in a conservation orchard with trees of different ages. For grapevine, one to five clones from a single individual per accession were phenotyped, and quantitative variables were averaged across clones – qualitative variables being the same across clones. For a given accession, the clones were planted in a single row, so there were no environmental replicates. Cabbage, grapevine, apple and foxtail millet were phenotyped in the field, while all other species were phenotyped in a greenhouse (Table S3). Note that sugar beet was phenotyped in controlled chamber where plant cycles were synchronized to ensure simultaneous flowering of wild (annual) and domestic (biennial) individuals. All specific phenotyping conditions are detailed in Table S3.

The number of traits measured within species varied widely, ranging from 11 traits in apple to 57 traits in sugar beet (Table S4 and S5), summing to 226 distinct traits across the 13 species (Table S5). A complete ontology of the traits can be found at https://doi.org/10.57745/QWEKVK. Traits were evaluated from germination to maturity and harvest. They included both common descriptors across most species, such as plant height or grain size, as well as species-specific traits, such as the acidity of grape must or the spiny or non-spiny leaves of eggplant (Table S4).

The traits measured fell into three categories (1) continuous quantitative variables, such as stem circumferences, pH and the Brix index (an estimate of the sugar content); (2) discrete quantitative variables, such as the number of internodes or seeds; (3) qualitative variables, such as flower colour or the relative position of styles within the flower. We used imaging software to assess some of the traits including the number of pollen grains in grapevine flowers (Gimenez et al., 2024) and several root traits in sugar beet (Bucksch et al., 2014).

### NIRS acquisition and analysis

We acquired Near InfraRed Spectra (NIRS) on leaves (NIRS_leaf_) for all species. In addition, NIRS were collected on seeds for eight of the 13 species (SNIRS1 and SNIRS2, Table S5), specifically those for which sufficient seed material was available. For NIRS phenotyping of leaves, we harvested three leaves or leaf sections per individual. Samples were oven-dried for 48 h at 40°C in coffee filters prior to measurement with a LabSpec 2500 NIR spectrometer. Drying has been shown not to significantly affect spectra (Borraz-Martínez et al., 2019). For each leaf, three spectra were collected, each obtained as the mean of 10 scans. Wavelengths ranged from 350 to 2500 nm at 1-nm intervals, yielding 2150 wavelengths per spectrum. For NIRS phenotyping on seeds (NIRS_seed_), we used a Perkin Elmer FT9700 NIR spectrometer. To account for sample heterogeneity within species, we performed four technical replicates per sample. Each spectrum was obtained as the mean of 32 scans, and wavelengths ranged from 700 to 2500 nm with a step of 8 cm^-1^, resulting on 1300 wavelengths.

We prepared the data as follows, using recommended procedures in Rincent et al. (2018) to control for additive and multiplicative effect of wavelength: (1) we loaded the spectra using the *asdreader* R package (https://github.com/pierreroudier/asdreader); (2) we discarded technical replicates from one individual whose deviation from the individual mean exceeded twice the standard deviation over at least 30% of the spectrum – apparent outlier replicates were also discarded; (3) we further normalized the spectra by sample, and computed their first derivative using a Savitsky-Golay filter (Savitzky & Golay, 1964) as implemented in the *signal* R package (https://cran.r-project.org/web/packages/signal/index.html) using a window size of 37 points and a filter order of two.

Because the wavelengths are numerous and not independent, we computed Principal Coordinate Analyses (PCoA) for leaf and seed samples separately. We used the projections of the individuals on the first two PCoA axes as two new independent traits. We evaluated whether clustering obtained with NIRS effectively grouped the technical replicates of the same individual. In order to do so, we conducted supervised hierarchical clustering of *g* groups, where *g* represents the number of individuals. The grouping of technical replicates was validated using the R package *flexclust* (Leisch, 2006) and the Rand index (Rand, 1971). The Rand index, which measures clustering similarity, ranges from zero to one, with the latter indicating perfect match between the observed and that predicted from individuals.

### Domestication syndrome characterization

To describe the domestication syndrome in each species, we applied a trait-by-trait approach. For quantitative traits, we tested differences in mean between forms using a mixed linear model:

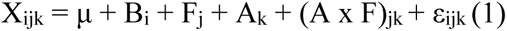

where X_ijk_ represents the phenotypic value of one trait in replicate i, in form j and accession k; μ is the phenotypic mean; B_i_ is the fixed effect of replicate i; F_j_ is the fixed effect of form j; A_k_ is the random effect of accession k, with A_k_ ∼ 𝒩(0, σ^2^_k_); (A x F)_jk_ is the random effect of the interaction between form j and accession k, with (A x F)_jk_ ∼ 𝒩(0, σ^2^_j_), σ^2^_j_ being the specific variance for form j; ε_ijk_ is the residual error, with ε_ijk_ ∼ 𝒩(0, σ^2^_E_).

When there was no replicate, we used a simple linear model:

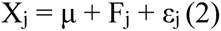

Where X_j_ represent the phenotypic value of a trait in form j; μ is the phenotypic mean; F_j_ is the fixed effect of form j; ε_j_ is the residual error, with ε_j_ ∼ 𝒩(0, σ^2^_E_). We used Student’s t-tests as implemented in the *lmertest* R package (Kuznetsova et al., 2017) to assess the significance of fixed effects. For qualitative traits we used Fisher exact tests to assess significant differences in the distribution. We calculated species-specific false discovery rate (FDR) q-values using the Benjamini–Hochberg procedure (Benjamini & Hochberg, 1995) based on the p-values for the form effect, applying a 5% threshold. We defined domestication-associated traits as those showing a significant main effect of form i.e., trait values that differed between domestic and wild forms. For graphical representation of trait values in each species, we constructed radarplot using the R packages *ggplot2* (Wickham, 2009) and *ggradar* (https://github.com/ricardo-bion/ggradar).

### Domestication syndrome convergence

To test for convergence, we assessed whether species shared more domestication-associated traits than expected by chance. First, we determined the observed number of domestication-associated traits unique to a species, or shared by two, or three or more species. Second, we applied a resampling procedure to generate a null distribution under the hypothesis of random sharing of domestication-associated traits (i.e., no convergence). To account for the structure of our data, we resampled traits using a design that mimicked both the number of traits measured per species and the pattern of trait sharing across species. Note that because we analyzed NIRS measured on seeds using the first two axes of a PCoA, with potentially different wavelengths contributing unequally to each axis across species; these axes are not homologous among species and were excluded from the convergence analysis. Of the remaining 341 measured traits across species (including traits shared by multiple species, Table S5), we identified *n* domestication-associated traits. Our procedure involved performing 10,000 resampling iterations in which *n* traits were randomly drawn from the *t* traits and designated as domestication-associated traits. For each resampling, we generated a simulated distribution that we compared to the observed values to compute p-values. Convergence was supported if the number of traits shared across three or more species exceeded that expected by chance, as determined by the simulated distribution.

### Computation of the multivariate phenotypic space

We aimed to compare the relative size of the multivariate phenotypic space between wild and domesticated forms across all species pairs. As the determinant of the variance-covariance matrix describing a space is related to its size, we used it as a measurement of the phenotypic space hypervolume. To compute the variance-covariance matrix in each species, we (1) normalized the quantitative traits across all individuals; (2) transformed qualitative traits by decomposing them into their constituent levels (e.g., a trait with five levels became five binary traits), scoring each as present or absent. Considering each form (wild and domesticated) within species, we further eliminated traits that were direct linear combinations of others. Finally, we computed the determinant of the variance-covariance matrix of traits for each form.

Since we removed linear combination of traits separately for wild and domesticated forms, the sizes of the matrices were not necessarily equal, making direct comparisons of determinants impossible. To address this, we normalized the determinants by taking the *r*-root, where *r* is the highest dimension between the wild and domestic matrices. We then calculated the log ratio of the size of the multivariate domesticated phenotypic space to that of the wild multivariate phenotypic space. A ratio greater than one indicated that domesticated forms explore a broader phenotypic space than their wild counterparts, while a ratio smaller than one indicated the reverse. We tested the influence of the number of samples and the number of traits on the hypervolume size by using Kendall and Spearman correlation tests. We used the same tests to assess correlations between the log ratio of the phenotypic spaces and the domestication timing in generations (Table S1).

### Computation of phenotypic spaces disjunction

We measured the extent of disjunction between the phenotypic spaces of wild and domesticated forms using the Pillai trace (Pillai, 1955) which quantifies how well grouping factors separate observations in multivariate data. We first performed a Factorial Analysis of Mixed Data (FAMD) on each dataset; we further considered all resulting axes as new multivariate variables. Pillai trace was then calculated in a MANOVA using the form (wild or domestic) as the explanatory variable and the FAMD axes as response variables. The resulting value ranged from zero, indicating complete overlap of the spaces, to one, indicating complete disjunction between the two forms.

Our comparative study is based on interspecific sampling. Despite efforts to harmonize sampling criteria across species, inherent heterogeneity remains and may bias estimates of the disjunction between wild and domestic phenotypic spaces. Indeed, depending on the species, the sampled wild and domestic accessions may have diverged to varying degrees. Increased divergence may result, for example, from sampling domesticated forms farther away from the domestication center–and therefore having experienced stronger drift to the wild gene pool–or from sampling domesticated forms that exhibited higher level of improvement.

In order to control for these sampling effects, we hypothesized that NIRS_leaf_ primarily captured variation unrelated with selection during domestication. In other words, while some wavelengths may be shaped by domestication-related selection, we expected the full spectral range to provide a baseline reflecting drift and selective processes unrelated with domestication. Previous studies have indeed shown that a substantial proportion of the variability in NIRS is explained by genome-wide SNP markers, indicating that most of the spectrum can be explained (Rincent et al., 2018) and predicted (Robert et al., 2022) based on overall genetic similarity among genotypes. To confirm our hypothesis, we first compared the PCoA obtained from strongly selected organs during domestication (NIRS_seed_) to the one obtained from NIRS_leaf_. Next, we determined wild and domestic multivariate phenotypic spaces on NIRS_leaf_ using all the axes of the corresponding PCoA as variables, and correlated the log ratio of domestic to wild space size with the corresponding ratio of genomic diversity taken from literature (Table S6) using Spearman and Kendall correlation tests. The latter was used as a proxy for drift.

By computing Pillai trace on the two first axes of PCoA computed on NIRS_leaf_, we further estimated the baseline disjunction (*Pillai_control_*) that we compared with values obtained for the Pillai trace computed from all other traits. We then corrected the disjunction of other traits by subtracting *Pillai_control_* to obtain a corrected space disjunction. In the following, we refer to the resulting index as the multivariate Phenotypic Divergence Index (mPDI). We also computed the mPDI per unit of baseline disjunction, by dividing mPDI by *Pillai_control_*.

We assessed correlations between mPDI and archaeological estimates of the domestication timing in generations (Table S1) as well as between mPDI and the log ratio of the size of the multivariate domesticated phenotypic space to that of the wild multivariate phenotypic space using Kendall and Spearman correlation tests; and tested the effect of mating system on the mPDI using a Student t-test.

### Similarity of trait matrices between wild and domesticated forms

For each form, we calculated the mean value of each trait by averaging across replicates (when multiple replicates were available, Table S3). We used two approaches to assess the similarity of trait matrices between wild and domesticated forms. In the first approach, we computed all pairwise phenotypic correlations between traits within each form using Spearman correlation coefficients. Note that we did not include a random accession effect when computing correlations. We next compared the absolute mean values of correlations between forms using a Student t-test. In the second approach, we computed phenotypic variance-covariance matrices and assessed their similarity between forms using the random skewers method (Cheverud & Marroig, 2007) as implemented in the *EvolQG* R package (Melo et al., 2016). This method is based on Lande’s equation (Lande, 1979), and evaluates how similarly two matrices respond to deformation. In short, each matrix is multiplied by one random scaled deforming vector, producing two matrices whose deformation can be described by a response vector. The correlation between the two response vectors is measured. This process is repeated a large number of times (10,000 in our case), yielding the average response vector correlation. The advantage of the random skewers method over a Mantel test, is that it provides a quantitative measure of the similarity between the variance-covariance matrices of wild and domestic traits, enabling comparisons across species. We assessed correlations between the similarity of the matrices and archaeological estimates of domestication timing in generations (Table S1) and mPDI with Kendall and Spearman correlation tests.

## Results

We measured 226 traits across 13 species, 185 of which were species-specific while 41 were shared by 2 to 11 species (Table S5). Such a diversity of species and traits poses a statistical challenge for performing comparisons across species. In addition to trait-by-trait analysis within species, we therefore developed a unifying statistical framework that includes the establishment of a multivariate phenotypic divergence index (mPDI) and the use of specific tools to compare the evolution of phenotypic spaces and trait correlations across species.

### Domestication syndrome

We characterized the phenotypic domestication syndrome within species. Domestication-associated traits were identified based on statistically significant differences in trait values between wild and domesticated forms. For example, in foxtail millet, we measured 15 traits, 12 of which were domestication-associated traits that differentiated wild and domesticated forms (Fig. 3a). Across systems, the percent of domestication-associated traits among all traits ranged from 26% in melon and 35% in grapevine to 81% in maize and 89.7% in common bean (Table S4; Figs S1-12).

**Figure 3:**
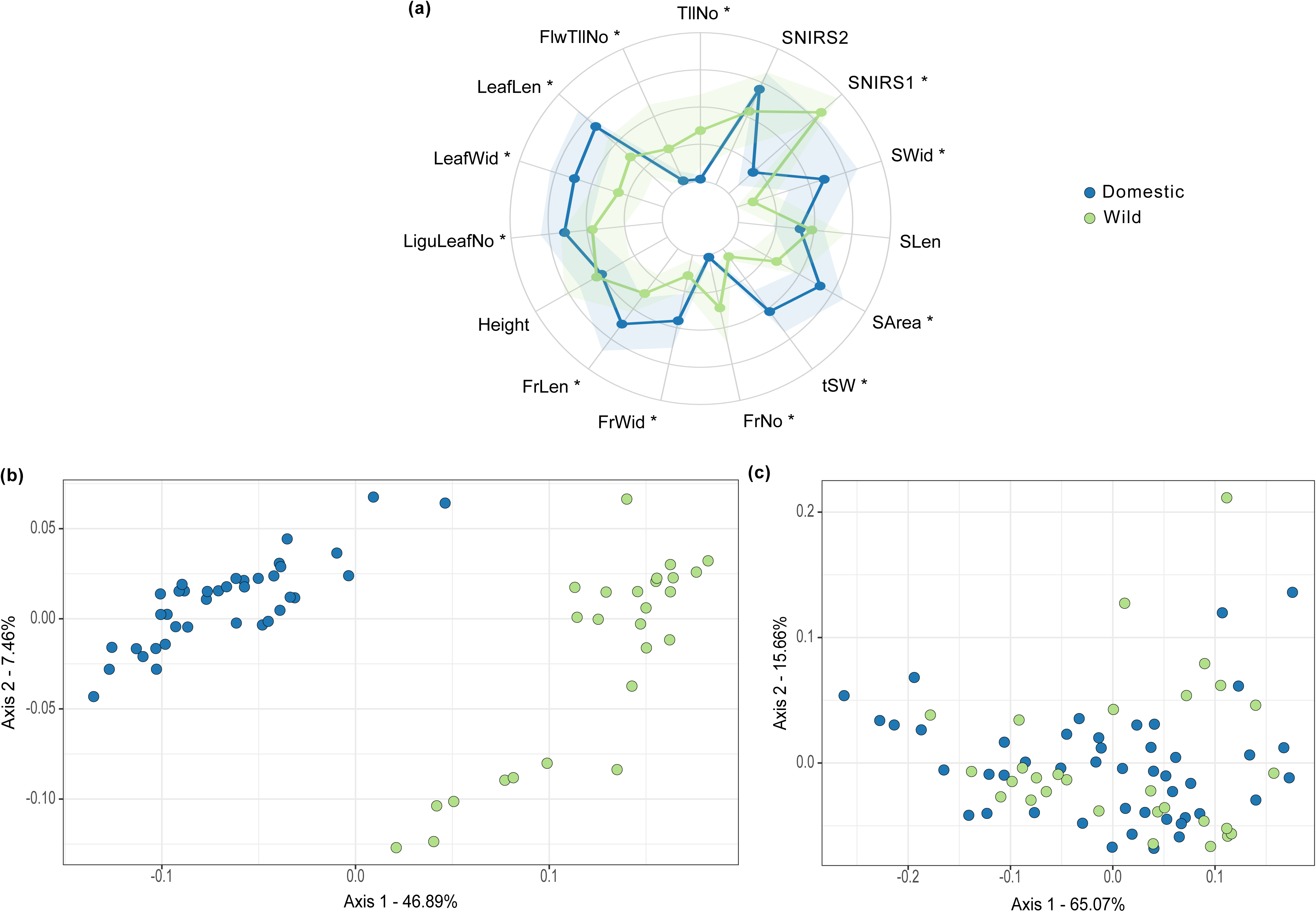
Domestication syndrome in foxtail millet. Spider-plot for quantitative traits with significance (*) between wild and domesticated forms at an FDR of 5% (a). First two axes of the PCoA computed from NIRS_seed_ (b) and NIRS_leaf_ (c). In (a) the dots represent the mean values, with the shaded area showing the standard error. TllNo: Tiller number per plant; SNIRS1: First axis of PCoA on NIRS_seed_; SNIRS2: Second axis of PCoA on NIRS_seed_; SWid: Seed width; SLen: Seed length; SArea: Seed area; tSW: Thousand seed weight; FrNo: Number of spikes per plant; FrWid: Spike width; FrLen: Spike length; Heigth: Plant heigth; LiguLeafNo: Number of liguled leaf; LeafWid: Leaf width; LeafLen: Leaf length; FlwTllNo: Flowering tiller number per plant.

In addition, as illustrated in foxtail millet for 8 out of 13 domestication-associated traits (Fig. 3a), the domesticated forms tended to display greater values than the wild forms, e.g., larger leaves and spikes. This pattern was repeatedly observed across species extending to domestication-associated traits related to plant and organ (leaf/spike/fruit/seed) size (Fig. 3a, Figs S1-S12). We also found a number of qualitative domestication-associated traits such as in common bean, where the colour, pattern and tendency for shattering in pods differed between wild and domesticated forms (Fig. S4, FrCol, FrPat and FrTension). Our findings also highlighted new interesting domestication-associated traits, such as the increase of petal length in eggplant (PtILen, Fig. S5) and the decrease in the number of secondary stems in common bean (NodeNo2, Fig. S4). We investigated seed/grain properties using NIRS, hypothesizing that domestication may have influenced their composition. Interestingly, we found significant differences in NIRS profiles between wild and domesticated forms (either on the first axis of the PCoA – SNIRS1, or the second – SNIRS 2 or both) in melon, einkorn wheat, foxtail millet, cabbage, maize, African rice, common bean and pearl millet (Fig. 2b, Fig. 3b, Figs S1-12).

### Convergence of domestication-associated traits

Among 78 species pairs, we found that 82% had at least one shared domestication-associated trait (Fig. S13). The average number of pairwise shared traits across all species was 2.6 and ranged from 0 – between African rice and eggplant or African rice and grapevine and cabbage, to 7 – between maize and tomato or foxtail millet and maize (Fig. S13). We formally tested phenotypic convergence for domestication-associated traits. While the number of unique domestication-associated traits to a given species did not significantly differ from expectation, we found a significant enrichment of domestication-associated traits (convergence) shared by three or more species (P-value=0.014, Fig. S14) indicating convergence. Conversely, this was accompanied by a deficit of traits shared by exactly two species (P-value=0.014, Fig. S14), consistent with the fixed total number of traits—where enrichment in one category entails a corresponding reduction in another.

Our dataset included several *Poaceae* species (5 of the 13 species, Table S1), a family known for phenotypic convergence of domestication-associated traits (Glémin & Bataillon, 2009). To ensure that observed convergence was not driven by this sampling, we repeated the analysis including only one of the *Poaceae* species at a time. The results remained consistent (P-values<0.01) with an enrichment of traits shared by three or more species. Altogether, our results indicate convergence in trait selection at a broad evolutionary scale, spanning multiple botanical families, including both monocots and dicots (Fig. 2).

The domestication traits displaying the greatest convergence are leaf size, leaf width and seed area shared respectively by 8, 7 and 9 species (Fig. 2b, LeafLen, LeafWid and SArea). Notably the *Poaceae* family encompassed an important number of shared domestication-associated traits (Fig. 2b), including grain area and width (SArea, SWid), the first axis of the PCoA based on NIRS measured on seeds (SNIRS1) and the number of tillers (TllNo).

### Comparative analysis of the multivariate phenotypic spaces

Within each species, we calculated the log ratio of the determinant of the domesticated variance-covariance matrix over that of the wild matrix using all traits to quantify the relative expansion or contraction of multivariate phenotypic diversity in domesticated forms compared to wild forms. We verified that this ratio was neither influenced by the sampling size of the domestic (Spearman, P-value = 0.69; Kendall, P-value = 0.58) and the wild (Spearman, P-value = 0.69; Kendall, P-value = 0.58) forms, nor by the difference between the two (Spearman, P-value = 0.69; Kendall, P-value = 0.58).

Three out of the thirteen pairs —namely einkorn wheat, apple and maize— presented positive log ratio values, indicating a larger phenotypic space in domesticated forms (Fig. 4). In most of the pairs (10 out of 13), the domesticated forms exhibited smaller phenotypic spaces than their wild counterparts, indicating a tendency for shrinkage during domestication. When looking at the absolute values of the log ratio, the pairs covered a range from 0.07 in sugar beet, to 2.2 in grapevine that displayed the largest shrinkage of phenotypic space. The log ratio was neither affected by mating system (Wilcoxon-Mann-Whitney, P-value = 0.83), nor by the timing of domestication (Spearman, P-value = 0.69; Kendall, P-value = 0.58).

**Figure 4:**
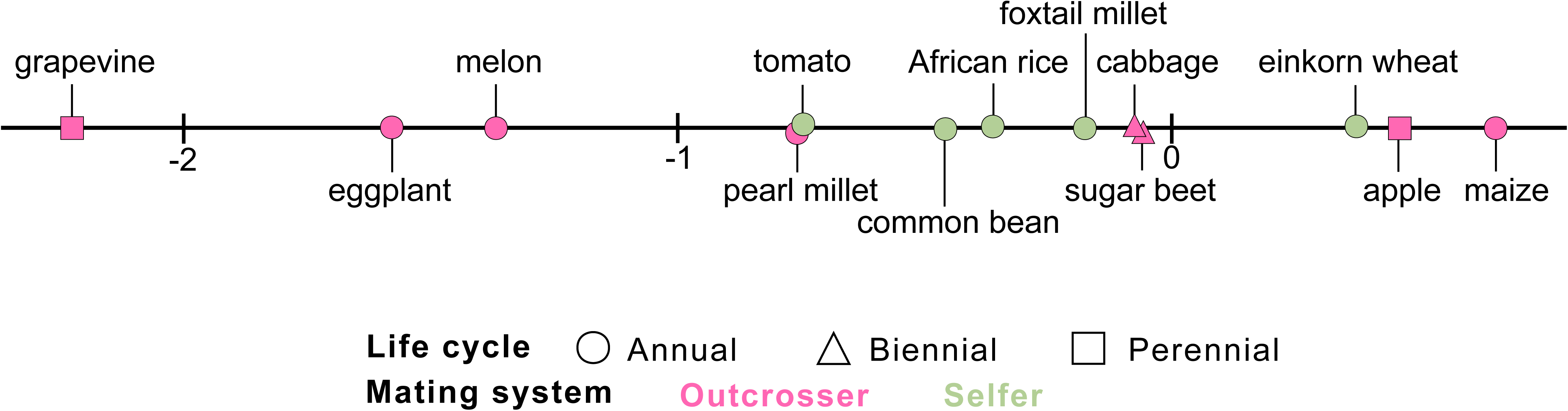
Relative size of domestic and wild multivariate phenotypic space. Log ratio of domestic over wild phenotypic space with positive/negative values indicating larger/smaller phenotypic space in domestic than in wild samples.

### Index of phenotypic divergence

Previous studies have shown that NIR spectra are able to discriminate between species (Lang et al., 2017; Meder et al., 2014) or varieties (Barbin et al., 2018), and can be used to capture genetic similarity between genotypes (Rincent et al., 2018). Reasoning that leaf chemical composition was less likely directly targeted by selection during domestication (except perhaps in leaf crops such as cabbage), we proposed to use NIRS profiling on leaves (NIRS_leaf_) as a control for inter-specific sampling biases to estimate the baseline disjunction between wild and domesticated forms. To verify this assumption, we first verified that NIRS_leaf_ captured a biological signal. Rand index values ranged from 0.92 in common bean to 0.96 in pearl millet confirming that NIRS_leaf_ effectively captured phenotypic diversity among individuals, with high consistency across replicates. Second, we verified that the values of the Pillai trace obtained from NIRS_leaf_ (Pillai_control_) were much lower than the ones computed on other traits (≥0.71 on other traits versus ≤0.16 for NIRS_leaf_ excluding apple, Table S6). Consistently, we found no distinction between wild and domesticated forms along the axes of the PCoA in contrast with what we obtained with NIRS profiling on seeds (SNIRS1 and SNIRS2) as illustrated in foxtail millet (Fig. 2b-c) and all other species (Figs S3-S12) except apple (Fig S2). In apple, Pillai_control_ was high (0.70, Table S6) reflecting our sampling of distant Asiatic wild accessions and European domestic accessions. We also verified that the mPDI values were not influenced by the number of traits measured in each species (Spearman: P-value= 0.57; Kendall: P-value=0.56). Finally, we computed the log ratio of phenotypic spaces using size of NIRS_leaf_ and found a positive correlation (Spearman, P-value = 0.04; Kendall, P-value = 0.04, Fig. S15) with the ratio of estimates of nucleotide diversity from the literature (Table S6) indicating that the evolution of the phenotypic space of NIRS_leaf_ seems mainly driven by genetic drift between wild and domesticated forms, further reinforcing its use as a control trait.

Based on these results, we treated NIRS_leaf_ as a control trait to compute a multivariate phenotypic divergence index (mPDI). Because the mPDI of apple was very close to zero (0.03, Table S6) and the disjunction of the phenotypic space was almost entirely due to the baseline divergence between forms, we excluded it from the rest of our analyses.

mPDI values ranged from 0.69 for sugar beet to 0.98 for melon with an average of 0.81 (Fig. 1b, Table S6). Note that mPDI values obtained for maize (0.77) were likely underestimated due to suboptimal greenhouse conditions (Table S3), which hindered data collection as many ears failed to produce seeds, resulting in poor characterization of the reproductive organs. All together our results indicated that wild and domesticated forms occupy distinct phenotypic spaces with some variation across species: overlap between phenotypic spaces was greater in sugar beet, tomato or pearl millet than in grapevine, common bean or melon (Fig. 1b). We found neither an effect of mating system on mPDI (Wilcoxon-Mann-Whitney: P-value= 0.76), nor a correlation between mPDI and the timing of domestication (Spearman: P-value= 0.64; Kendall: P-value= 0.58). We further tested the effect of mating system on the mPDI per unit of disjunction unrelated with domestication and found a tendency for selfers to be less disjoint than outcrossers (Fig. 1c, Wilcoxon-Mann-Whitney: P-value= 0.15).

### Comparative analysis of trait correlations

Common bean and tomato displayed significantly higher mean absolute correlations between traits than their wild counterparts. Conversely, cabbage, eggplant, einkorn wheat, foxtail millet, grapevine, melon and sugar beet displayed lower mean absolute correlation values in domesticated forms than in wild forms. In the other five wild/domestic pairs — African rice, apple, maize and pearl millet — we detected no difference in the average absolute correlation values between domestic and wild form (Table S7).

Random skewers method was used to compute the similarity between domestic and wild variance-covariance matrices of traits (Fig. 5). Similarity values ranged from 0.36 in tomato to 0.99 in grapevine, spanning a continuous gradient. We found no impact of the mating system (Wilcoxon-Mann-Whitney: P-value=0.09), and matrices similarity did not correlate with phenotypic disjunction (Spearman: P-value= 0.72; Kendall: P-value= 0.85). However, it did correlate negatively with domestication timing (Fig. 5; Spearman: P-value= 0.004; Kendall: P-value= 0.01).

**Figure 5:**
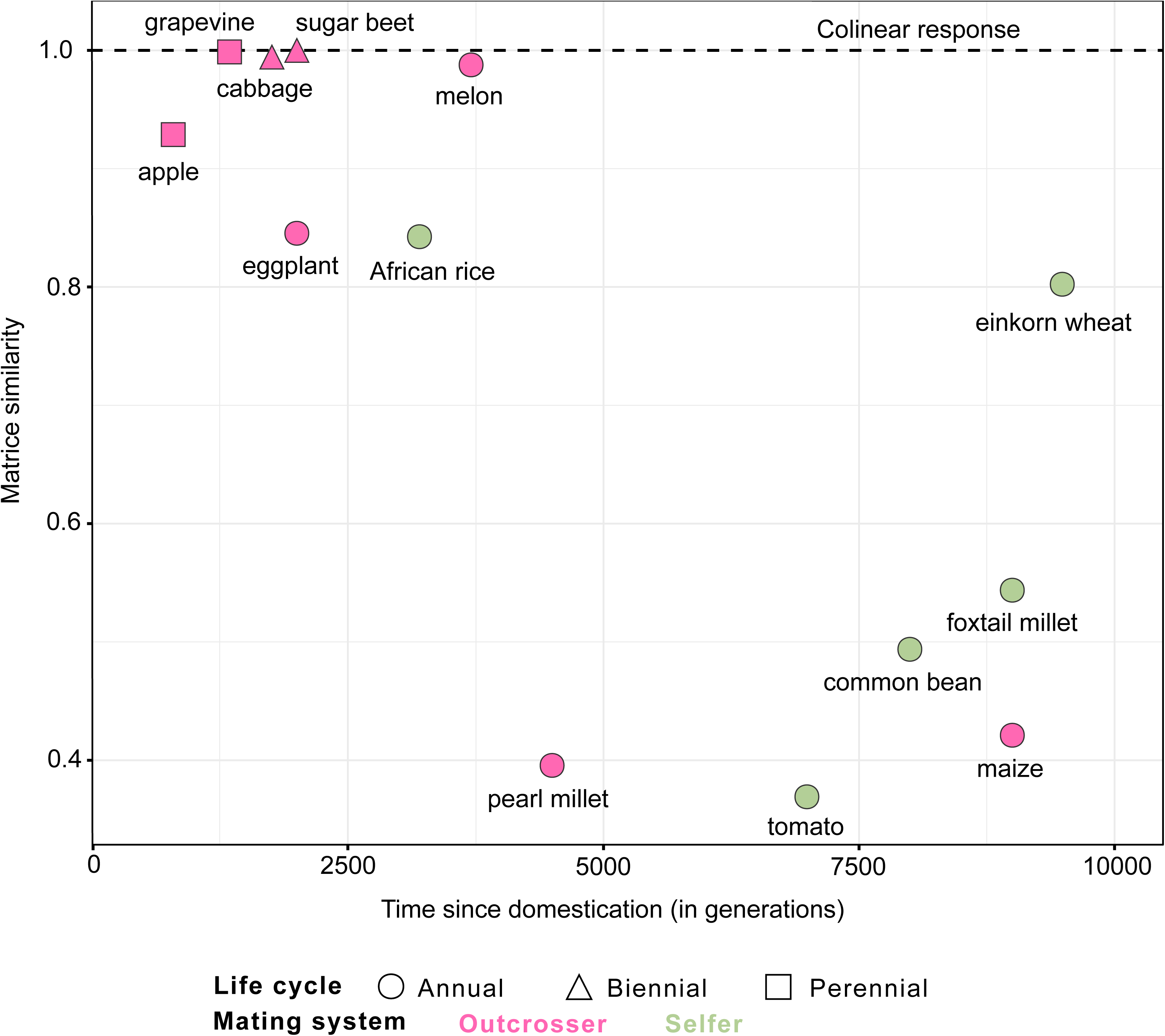
Correlation between similarity of wild and domestic variance-covariance matrices and domestication timing (Table S1). A matrix similarity of 1 indicates colinear responses to deformation.

## Discussion

Since Darwin (1868), domestication has served as an *in vivo* experiment for studying phenotypic evolution, offering replicated experiments to explore how divergent selection has shaped traits over time. However, most studies have focused within species, aiming to identify domestication traits and their underlying genetic changes to infer mechanisms of adaptation to human-made environments. Surprisingly, broader patterns of phenotypic shifts and the key parameters driving their evolution remain poorly understood.

A major limitation for cross-species comparisons arises from sampling biases. To limit these biases, we focused on accessions near the center of domestication to minimize genetic structuring and avoid confounding effects of recent breeding, with apple as the sole exception. We phenotyped these accessions using standardized measurements across a diverse set of traits, representing both species-specific and shared traits. We also customized a statistical framework for cross-species analyses. A key contribution is the use of NIRS_leaf_ as a low-cost, easily measurable control trait. Building on this, we established a multivariate Phenotypic Divergence Index (mPDI).

Evaluating traits across species in a single common garden was impractical due to differences in species geographical origins and life cycles. To minimize environmental effects, we established common gardens within each species (Table S3). Because each species required different cultivation conditions, such as photoperiod and temperature, those not suited to the local climate were evaluated in controlled greenhouse conditions. These controlled environments may have amplified or reduced differences between wild and domesticated forms—for example, differences in maize appear reduced— which is an important limitation of our study. For perennial or biennial species, the lack of replicates further limited our ability to control for environmental effects. Despite these caveats some clear trends emerged. The proposed framework is broadly applicable, does not require genomic data, and can be extended to additional species, accessions, and traits, enabling wider testing of the trends identified here.

### Convergence of domestication-associated traits

The convergence of domestication-associated traits has been well documented in cereals for traits such as seed shattering. It has revealed notable cases of underlying genetic convergence where orthologous genes contribute to variation of the same trait across different species (Patterson et al., 2023). Recently, studies have explored convergence over deep evolutionary timescales in crops from distantly related families, once again highlighting cases of genetic convergence. An example is the *G* gene, which influences seed dormancy variation in rice, tomato, soybean, sorghum and maize (Rendón-Anaya & Herrera-Estrella, 2018). At the phenotypic level, however, meta-analyses of convergence across species struggle with the diversity of experimental designs and disparity of measurement methods. By grouping domestication syndrome traits of 203 crops in nine classes associated to function, Meyer, DuVal, et al. (2012) have hinted convergence in some functions like the production of secondary metabolites (66% of crops). Nevertheless, the statistical significance of these observations was not tested and remains to be established. This necessitates a customized test, as the number of measurable shared traits between species decreases with increasing divergence across taxa. Here, by relying on a specific sampling strategy across species, and a resampling test procedure, we demonstrated that when domestication-associated traits were shared, they tended to be shared by more than two species (Figs. S13-S14).

Convergence was found for a diversity of traits. In *Poaceae*, for example, we formally established the convergence observed for reduced tillering, spike and seed size increase (Glémin & Bataillon, 2009). Convergence extended to more distant taxa like common bean or melon as previously reported (Meyer et al., 2012). We showed increases in leaf length and plant height across eight and seven species, respectively (LeafLen, Height, Fig. 2b), consistent with the well-documented trend of gigantism in crops (Milla & Matesanz, 2017). Noteworthy, seeds increased in size in nine species (Fig. 2b, SArea), a consequence of domestication known to extend to most crops. Indeed, seed enlargement during domestication is thought to have arisen initially as an indirect consequence of selection for deeper burial and increased interplant competition (Kluyver et al., 2013, 2017), and linked to the general increase of organ size by allometry (Milla & Matesanz, 2017). Thus, even though seeds are not always the consumed organ, convergence for this trait is found across taxa.

Interestingly, NIRS measured on seeds (SNIRS1, SNIRS2, Fig. 2b) was found to be a domestication-associated trait for all systems on which it was measured, even on systems where seeds were likely to have not been selected, e.g., cabbage and apple. For those crops, given that NIRS is a predictor of grain quality, it implies that the changes observed following domestication are not only relative to allometry but also to quality for a variety of species, as previously described by chemical composition analysis in wheat or maize (Fang et al., 2020; Zeibig et al., 2022). These changes may be related to the (partial) loss of seed dormancy during domestication, as it is linked to changes in seed chemical composition (Penfield, 2017; Prudente & Paiva, 2018).

### Evolution of the size of multivariate phenotypic spaces

Domestication has led to a loss of genomic diversity due to both demographic and selective processes (Yamasaki et al., 2005) that have reduced effective population sizes, thereby decreasing standing variation. One consequence of this loss is a depletion of genetic variance of traits under selection, as shown in maize (Yang et al., 2019), a pattern that extends to traits that are not directly targeted by selection (Rolhauser et al., 2024). Our results support the predicted loss of phenotypic variance, as evidenced by shrinkage of the multivariate phenotypic space in domesticated forms compared to their wild counterparts in 10 of the 13 species. However, we found no correlation between this ratio and the corresponding ratio of genomic diversity (Table S6; Spearman P-value=0.79; Kendall P-value=0.86), consistent with studies indicating that domestication affects only a small fraction of the genome—for example, 7.6% in maize (Hufford et al., 2012). Note that we found no effect of the mating system on the ratio of phenotypic diversity between domestic and wild forms, consistent with the hypothesis that selfing does not impair the response to directional selection (Noël et al., 2017).

### The multivariate phenotypic divergence index

In order to better understand the evolutionary mechanisms driving phenotypic divergence between wild and domesticated forms, it is essential to quantify this divergence using a measure that allows cross-species comparisons. A previous attempt proposed a semi-qualitative index, in which species were ranked according to their level of domestication from low (1) to high (5) (Dempewolf et al., 2008). Here, we propose a multivariate divergence quantitative index, the mPDI. Our results suggest that the mPDI is a reliable index. For instance, the common bean, which is known for its significant phenotypic disjunction (García et al., 1997; Peña-Valdivia et al., 2011) exhibits high mPDI (Fig. 1b). Conversely, sugar beet a very recent to domestication displays one of the lowest mPDI value (Fig. 1b). Note that in apple, mPDI was very close to zero, which can be explained by our sampling in this perennial species. Indeed, our domestic accessions originated from West Europe, whereas wild accessions were sampled in Central Asia. The Pillai value computed on all traits was thus almost equal to the value of the Pillai_control_ obtained with NIRS_leaf_, in line with the drift observed during apple expansion to the west (Cornille et al., 2014).

Overall, we found a high disjunction of phenotypic spaces between wild and domesticated forms across all species except apple (mPDI ≥ 0.69, Fig. 1b). Previous studies focusing on two traits had proposed rather contradictory hypotheses, suggesting either a disjunction (Glémin & Bataillon, 2009) or that domesticated forms represented a subset of the wild phenotypic space (Milla et al., 2015). Our characterization, extended to multivariate spaces, clearly supports the first hypothesis, confirming an expected outcome of divergent selection.

Phenotypic divergence is influenced by both genetic drift and selection. We therefore hypothesized that if it was primarily driven by drift, phenotypic divergence should increase with the time since domestication. While one of the youngest domesticates, sugar beet, and one of the oldest domesticate, common bean, were indeed found at the opposite extremes of mPDI values (Fig. 1b), we did not find a significant trend. This suggests that selection dominates the observed pattern. This conclusion is supported by the lack of a relationship between the phenotypic divergence measured in our study and estimates of genomic differentiation between wild and domesticated forms, estimated from previous studies (Table S6). Note that phenotypic divergence also reflects traits evolvability as measured by standing genetic variance (Opedal et al., 2023). We are thus capturing multiple effects with the mPDI, which are hard to decipher. Because selfers can adapt faster than outcrossers (Ronfort & Glemin, 2013) potentially accelerating phenotypic divergence, we tested whether the mating system had an effect on the mPDI. We found no effect, and when the mPDI was expressed per unit of disjunction estimated from the control trait, we observed the opposite trend, namely that selfers tended to fall in the lower range of values (Fig. 1c). However, this result should be interpreted with caution due to the binary classification of species as either selfers or outcrossers. In practice, many selfing crops exhibit low levels of outcrossing, which can be sufficient to mix alleles and phenotypes (Morjan & Rieseberg, 2004). Conversely, in outcrossing species, barriers to gene flow between wild and domesticates exist–such as the *GA1* and *GA2* loci that prevent maize pollen from fertilizing teosintes (Chen et al., 2022; Evans & Kermicle, 2001) among other examples. Quantitative variation in the intensity of gene flow, rather than the mating system *per se*, may play a more critical role in determining mPDI rankings, but gene flow remains challenging to quantify.

As a measure of phenotypic divergence between wild and domesticated forms, and thus of adaptation to anthropogenetic niches, mPDI could be correlated with the fitness of these forms in human-altered habitats or with their occurrence across environments differently affected by human activity. Following the domestication framework proposed by Lord et al. (2025), species with higher mPDI values are expected to be better adapted to human-mediated environments. Unlike other indices of phenotypic divergence, such as Darwin’s (Haldane, 1949), Haldane’s (Gingerich, 1993) or Qst (Spitze, 1993), mPDI offers a multivariate quantitative measure that allows cross-species comparisons and can be readily applied beyond domestication contexts—for instance, to assess local adaptation in populations (Fustier et al., 2019) or to study phenotypic evolution in experimental evolution assays (Desbiez-Piat et al., 2023). Note that while easily applicable in distantly related plant taxa, the use of NIRS on specific homologous organ as a control trait may be impractical for animal species.

### Reorchestration of correlations between traits

The concept of domestication syndrome not only refers to a series of traits that are different between forms, but also to the correlations between them (Milla et al., 2015). The impact of phenotypic shifts across multiple traits on the evolution of phenotypic spaces is still not fully understood. Phenotypic evolution follows a genetic line of least resistance corresponding to traits with high genetic variance (Mallard, Afonso, et al., 2023; Opedal et al., 2023). The rate of this evolution is determined by the alignment between this line and the direction of selection (Mallard, Afonso, et al., 2023), and therefore on the correlation between traits. Classically, a genetic matrix of traits correlations (G-matrix) is computed. Here we relied on the phenotypic matrix of traits correlations as a proxy for the G-matrix (Cheverud, 1988).

We found that traits correlations changed during domestication, although with no clear pattern for increase or decrease across species. Previous studies predicted an increase in genetic correlations between traits in domestic compared to wild forms (Burban et al., 2022), notably because of the loss of genetic diversity in the domestic compartment and pleiotropic effect of genes, e.g. *GL4* in African rice for reduced grain size and non-shattering (Wu et al., 2017) and *TB1* in maize for modification of plant and root architecture and spikelet development (Stitzer & Ross-Ibarra, 2018). However, traits like seed shattering and grain size have been shown to evolve concomitantly but at different rate in cereals (Fuller et al., 2012). Domestication have also led to a decoupling of certain functions, such as root and leaf resource acquisition (Roucou et al., 2018).

Interestingly, we showed that the more time has passed the greater the uncoupling of variance-covariance matrices, as indicated by the loss of similarity between wild and domestic matrices (Fig. 5). This suggests that a general hallmark of domestication may involve a reorganization of trait correlations, a pattern previously identified in different species (various herbaceous crops, Milla et al., 2014; wheat, Roucou et al., 2018; maize, Yang et al., 2019). As time passes, the interplay of mutation and selection reshapes the structure of the G-matrix (Dugand et al., 2021; Houle et al., 2017; Mallard, Noble, et al., 2023), potentially leading to an alignment to the adaptative landscape (Hohenlohe & Arnold, 2008; Hunt et al., 2007). Consequently, as time since domestication increases, the G-matrices of wild and domesticated forms may diverge as they become aligned with their respective axes of selection. Altogether, these observations echoes findings that recombination tends to increase during domestication (Ross-Ibarra, 2004) or, at minimum, that the recombination landscape is reshaped along the genome (Bursell et al., 2025). Such novel recombination dynamics can influence phenotypic variation (Pan et al., 2016; Wagner, 2011) by breaking correlation between traits, potentially accelerating G-matrix alignment with the direction of selection in the anthropogenic niche.

## Conclusion

In this study, we present the first comparative analysis of plant domestication across multiple species. We introduce a novel phenotypic divergence index (mPDI), that does not require genomic data and is applicable across all domesticated species, thereby providing a standardized framework for cross-species comparisons. By characterizing the domestication syndrome in 13 species, we demonstrated that near-infrared spectroscopy (NIRS) measurements on grains contribute to the domestication syndrome in many species, highlighting the potential of the NIRS technology for both calculating the mPDI and characterizing domestication-related changes.

Our findings support several key hypotheses of domestication: trait convergence, disjunction between wild and domestic phenotypes, a relative reduction of the phenotypic space in domesticated forms, and a reorganization of trait correlations. Importantly, we reveal the emergence of shared patterns across species, independent of domestication timing or life-history traits, indicating the existence of general principles underlying plant domestication.

## Acknowledgments

We are very grateful to Roberto Papa and Elena Bitocchi (Università Politecnica delle Marche, Ancona, Italy) for providing common bean seeds; Alain Label-Richardson (Université Rennes, France), Lorenzo Maggioni (European Cooperative Program for Plant Genetic Resources, Roma, Italy), Roland Von Bothmer (Swedish University of Agricultural Sciences, Alnarp, Sweden), and Charlotte Allender & Nick Fenby (University of Warwick, UK) for providing cabbage seeds; Monika Höfer (Julius Kühn Institute - Federal Research Institute for Cultivated Plants) for managing apple genetic ressource and to Amandine Cornille (NYU Abu Dhabi) for discussing the sampling design of apple trees. We thank the technical staff Virginie Heraudet, Nathalie Galic, Cécilia Zokly, Arnaud Leclerc, Victoria Rozon and Florie Le Gentil of IDEEV greenhouse for the maintenance of plants. We thank the experimental unit A2M for the maintenance of plants and the CRB-Lèg for managing the vegetable genetic resources at INRAE Avignon. We also thank Yaël Bourgain, Dominique Emmanuel, Djouheina Saouli, Laura Courreyan, Martin Simonoviez, Lucie Guerrin, Violette van der Horst and Theophile Thomas for their contribution to data collection. We thank Leila Zekraoui at IRD for her contribution to the discussion on the phenotyping strategy. We thank Martin Ecarnot at INRAE for the access and support at the NIRS plateform ARCAD in Montpellier. We thank Betty Gimenez, Odile Peyron, Sébastien Joannin and Laurent Bouby for the counting of grapevine pollen grains. We thank Alain Ghesquière for the photography of African rice (© IRD). We also deeply thank Mark Chapman and two anonymous reviewers for their insightful comments on the manuscript. The authors used ChatGPT exclusively for language editing and not for generating any content. This work was supported by Agence Nationale de la Recherche (ANR-19-CE32-0009-02). A.W. was financed for 18 months by a doctoral contract from Institut National de Recherche pour l’Agriculture, l’Alimentation et l’Environnement (INRAE). GQE-Le Moulon benefits from the support of Saclay Plant Sciences-SPS (ANR-17-EUR-0007) as well as from the Institut Diversité, Ecologie et Evolution du Vivant (IDEEV).

## Author contributions

SG, YV, CD, MIT designed the research. AW, HB, SG, KA, PRG, DM, YV, CD, MIT set-up the experiments. KH, CM, TL, SG, KA, PRG, DM, YV, CD, MIT designed the sampling. AW, HB, AR, MB, MB, PS, AD, AP, KH, CM, TL, KA, PRG, DM, YV, CD, MIT performed the experiments and collected the data. AW, RR, SG, DM, YV, MIT designed the analysis pipeline. AW performed all analyses, and prepared all figures and tables. AW, HB, RR, CM, TL, SG, KA, PRG, DM, YV, CD, MIT contributed to data interpretation. MIT coordinated the project. AW and MIT wrote the manuscript. SG, YV, CD, PRG, DM provided comments on the manuscript.

## Data availability

Phenotypic data and NIR spectra are available on https://doi.org/10.57745/QWEKVK. R scripts are available on the INRAE Forge at https://forge.inrae.fr/gqe-gevad/domisol_phenotypic_spaces. Ontology of measured traits at the MIAPPE format is available at https://doi.org/10.57745/QWEKVK.

## Supporting Information

Figure S1: Domestication syndrome in African rice. Spider-plot for quantitative traits with significance (*) between wild and domesticated forms at an FDR of 5% (a). Cleveland plot of qualitative traits (b). First two axes of the PCoA computed from NIRS_seed_ (c) and NIRS_leaf_ (d). In (a) the dots represent the mean values, with the shaded area showing the standard error. Abbreviation meaning of traits can be found in table S4.

Figure S2: Domestication syndrome in apple. Spider-plot for quantitative traits with significance (*) between wild and domesticated forms at an FDR of 5% (a). First two axes of the PCoA computed from NIRS_leaf_ (b). In (a) the dots represent the mean values and the shaded areas denote standard error. Abbreviation meaning of traits can be found in table S4.

Figure S3: Domestication syndrome in cabbage. Spider-plot for quantitative traits with significance (*) between wild and domesticated forms at an FDR of 5% (a). Cleveland plot of qualitative traits (b). First two axes of the PCoA computed from NIRS_seed_ (c) and NIRS_leaf_ (d). In (a), the dots represent the mean values and the shaded areas denote standard error. Abbreviation meaning of traits can be found in table S4.

Figure S4: Domestication syndrome in common bean. Spider-plot for quantitative traits with significance (*) between wild and domesticated forms at an FDR of 5% (a). Cleveland plot of qualitative traits (b). First two axes of the PCoA computed from NIRS_seed_ (c) and NIRS_leaf_ (d). In (a), the dots represent the mean values and the shaded areas denote standard error. Abbreviation meaning of traits can be found in table S4.

Figure S5: Domestication syndrome in eggplant. Spider-plot for quantitative traits with significance (*) between wild and domesticated forms at an FDR of 5% (a). Cleveland plot of qualitative traits (b). First two axes of the PCoA computed from NIRS_leaf_ (c). In (a), the dots represent the mean values and the shaded areas denote standard error. Abbreviation meaning of traits can be found in table S4.

Figure S6: Domestication syndrome in einkorn wheat. Spider-plot for quantitative traits with significance (*) between wild and domesticated forms at an FDR of 5% (a). First two axes of the PCoA computed from NIRS_seed_ (b) and NIRS_leaf_ (c). In (a), the dots represent the mean values and the shaded areas denote standard error. Abbreviation meaning of traits can be found in table S4.

Figure S7: Domestication syndrome in grapevine. Spider-plot for quantitative traits with significance (*) between wild and domesticated forms at an FDR of 5% (a). Cleveland plot of qualitative traits (b). First two axes of the PCoA computed from NIRS_leaf_ (c). In (a), the dots represent the mean values and the shaded areas denote standard error. Abbreviation meaning of traits can be found in table S4.

Figure S8: Domestication syndrome in melon. Spider-plot for quantitative traits with significance (*) between wild and domesticated forms at an FDR of 5% (a). Cleveland plot of qualitative traits (b). First two axes of the PCoA computed from NIRS_seed_ (c) and NIRS_leaf_ (d). In (a), the dots represent the mean values and the shaded areas denote standard error. Abbreviation meaning of traits can be found in table S4.

Figure S9: Domestication syndrome in maize. Spider-plot for quantitative traits with significance (*) between wild and domesticated forms at an FDR of 5% (a). Cleveland plot of qualitative traits (b). First two axes of the PCoA computed from NIRS_seed_ (c) and NIRS_leaf_ (d). In (a), the dots represent the mean values and the shaded areas denote standard error. Abbreviation meaning of traits can be found in table S4.

Figure S10: Domestication syndrome in pearl millet. Spider-plot for quantitative traits with significance (*) between wild and domesticated forms at an FDR of 5% (a). Cleveland plot of qualitative traits (b). First two axes of the PCoA computed from NIRS_seed_ (c) and NIRS_leaf_ (d). In (a), the dots represent the mean values and the shaded areas denote standard error. Abbreviation meaning of traits can be found in table S4.

Figure S11: Domestication syndrome in sugar beet. Spider-plot for quantitative traits with significance (*) between wild and domesticated forms at an FDR of 5% (a). Cleveland plot of qualitative traits (b). First two axes of the PCoA computed from NIRS_leaf_ (c). In (a), the dots represent the mean values and the shaded areas denote standard error. Abbreviation meaning of traits can be found in table S4.

Figure S12: Domestication syndrome in tomato. Spider-plot for quantitative traits with significance (*) between wild and domesticated forms at an FDR of 5% (a). Cleveland plot of qualitative traits (b). First two axes of the PCoA computed from NIRS_leaf_ (c). In (a), the dots represent the mean values and the shaded areas denote standard error. Abbreviation meaning of traits can be found in table S4.

Figure S13: Convergence in domestication-associated traits. The number of traits measured per species and shared between species is shown (a) along with the subset of domestication-associated traits (b).

Figure S14: Testing for convergence of domestication-associated traits. Observed number of domestication traits either unique or shared by two, or three or more species are shown with the expected number of shared domestication traits as established by a resampling procedure. Significance was tested by Student t-test (P-values are indicated when significant; * P < 0.05; ** P <0.01).

Figure S15: Correlation between the log ratio of domestic to wild NIRS_leaf_ phenotypic space size with the ratio of domestic to wild genomic diversity (values from the literature, Table S6).

Table S1: Description of the 13 pairs of species with estimate of the domestication timing, domestication center used in this study, mating system and life cycle.

Table S2: Passport data of sampled accessions.

Table S3: Number of traits measured and growing conditions during phenotyping in each species.

Table S4: List of measured traits for each species, and domestication-associated (DA) traits with corresponding q-values and percentage of DA-traits in each species.

Table S5: Number of traits measured per species (columns) and number of species for which a given trait was measured (shared traits in rows). Traits are described Table S4.

Table S6: Table S6: Ratio of domestic (D) over wild (W) multivariate phenotypic space, mDPI, Pillai trace computed on all traits and on the control trait, and estimates from the literature of the ratio (D/W) of genomic diversity and Fst.

Table S7: Average absolute pairwise correlation between traits, computed from all pairwise correlations in wild and domesticated forms. P-values from Student t-tests.

