## Supplementary figures and images for "Evolution of crop phenotypic spaces through domestication"

### Figure S1

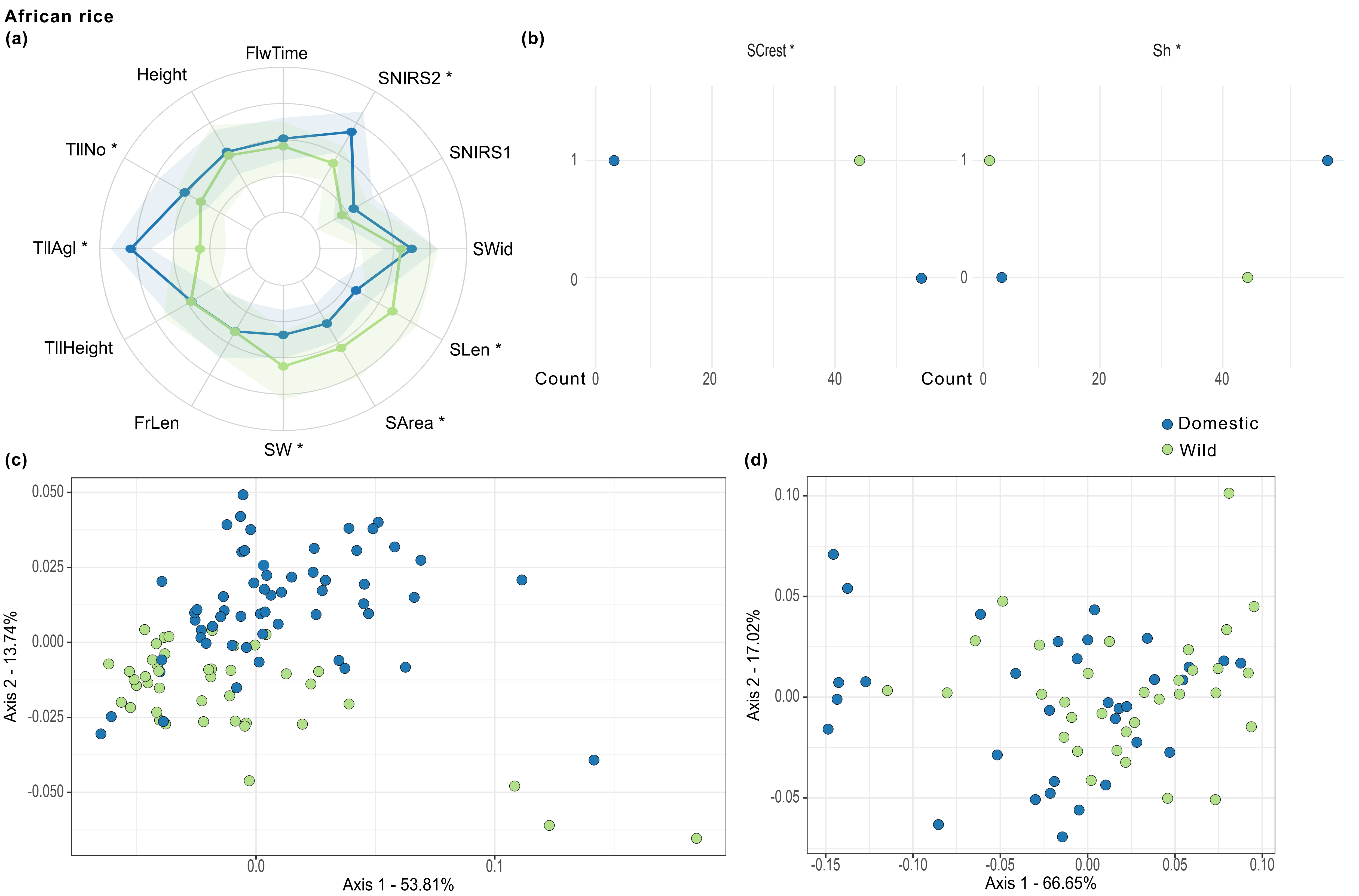

### Figure S2

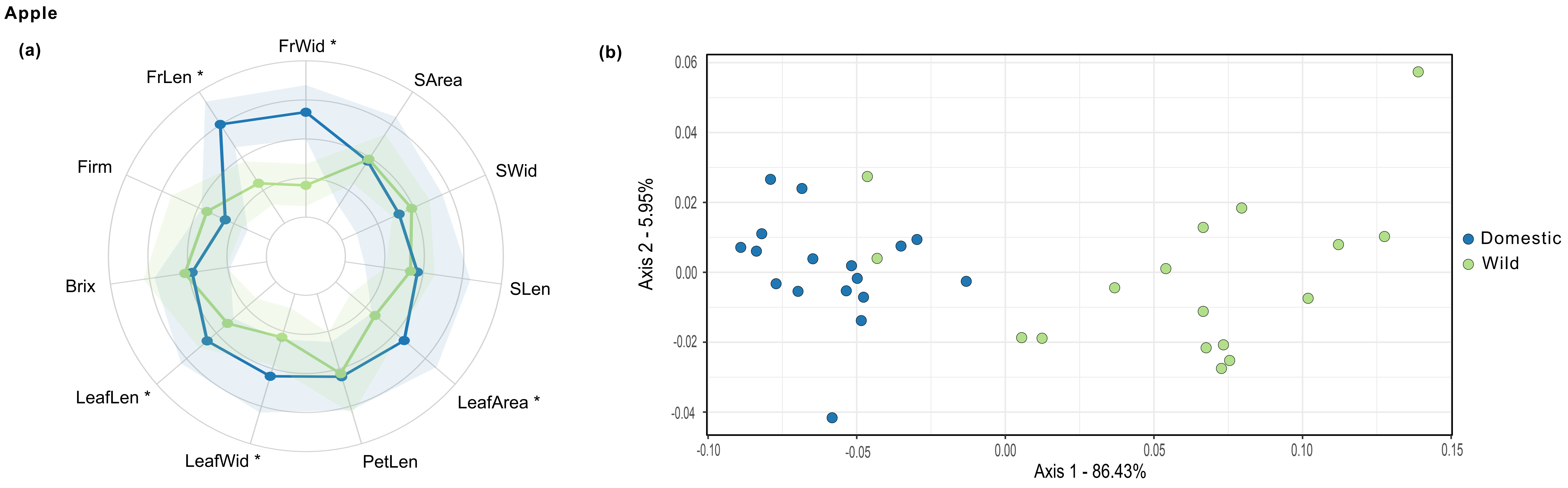

### Figure S3

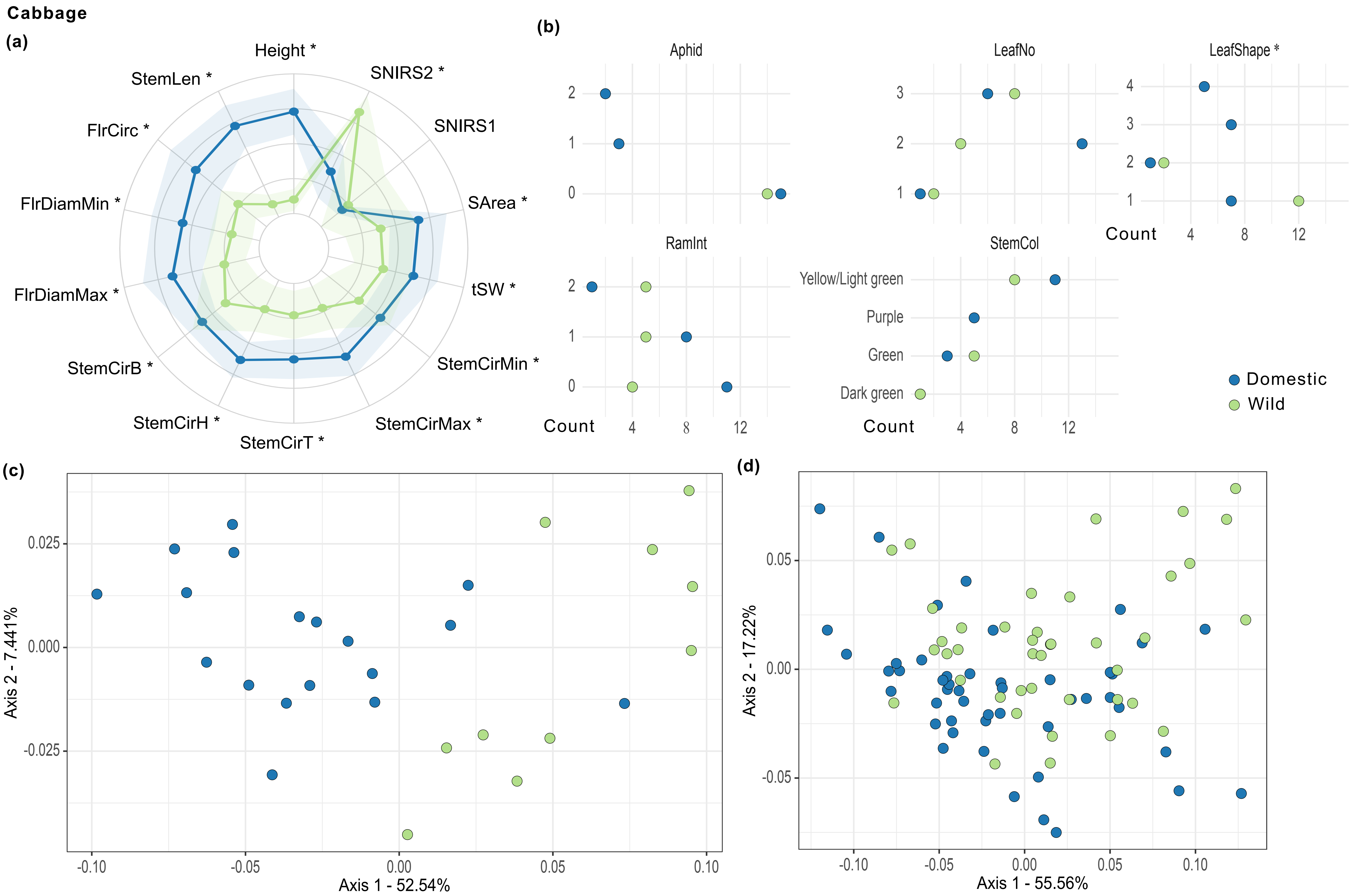

### Figure S4

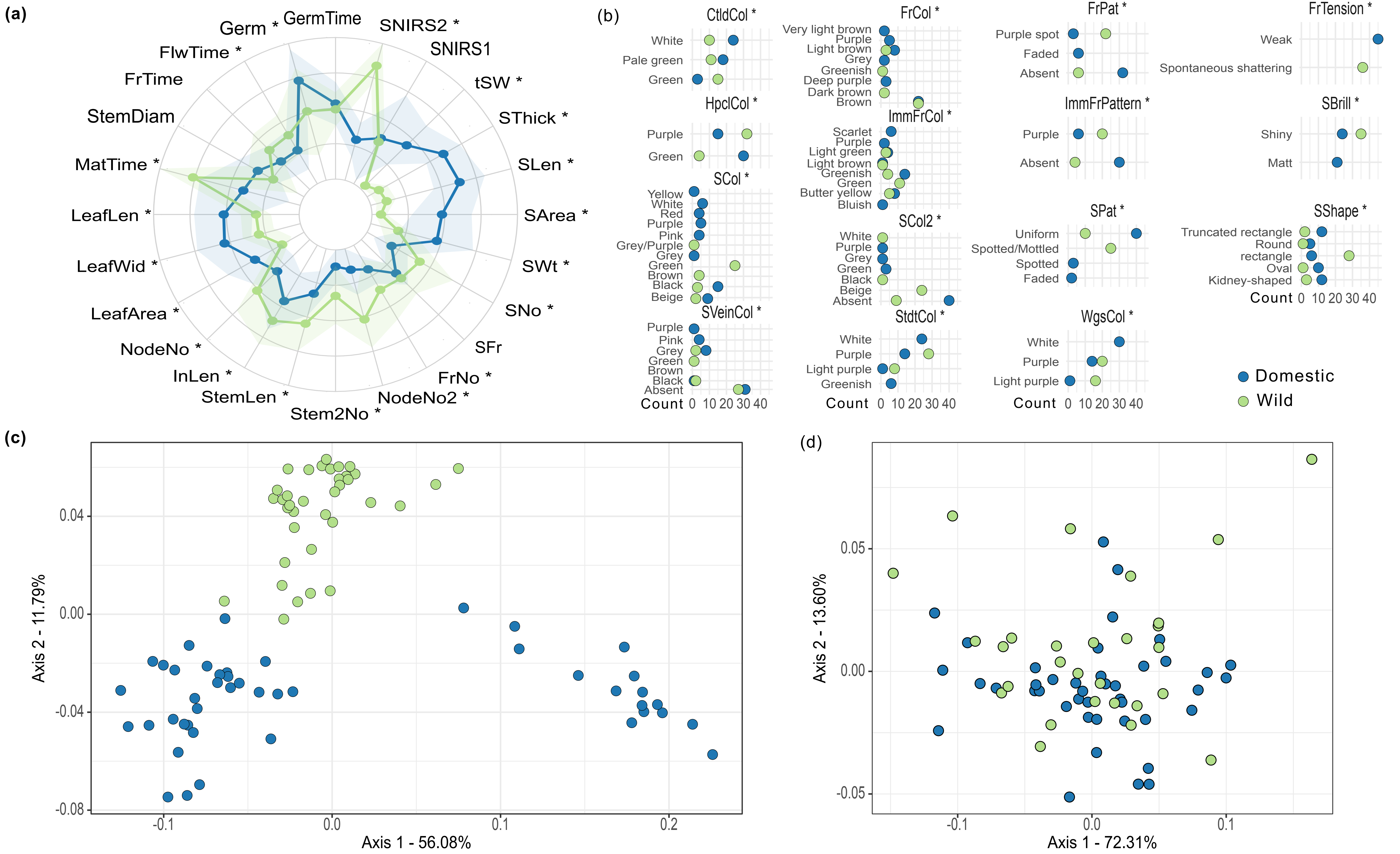

### Figure S5

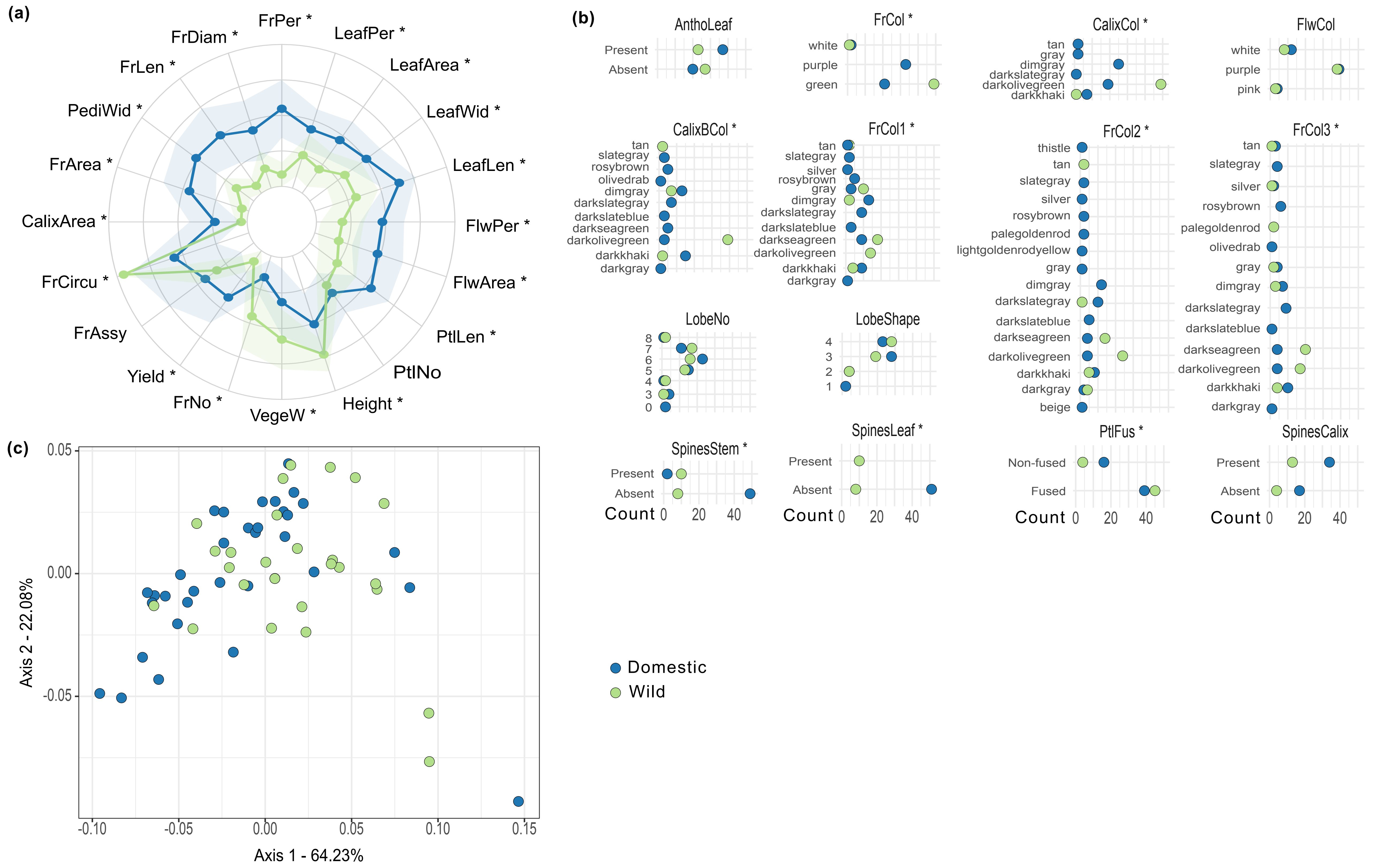

### Figure S6

**Einkorn wheat**

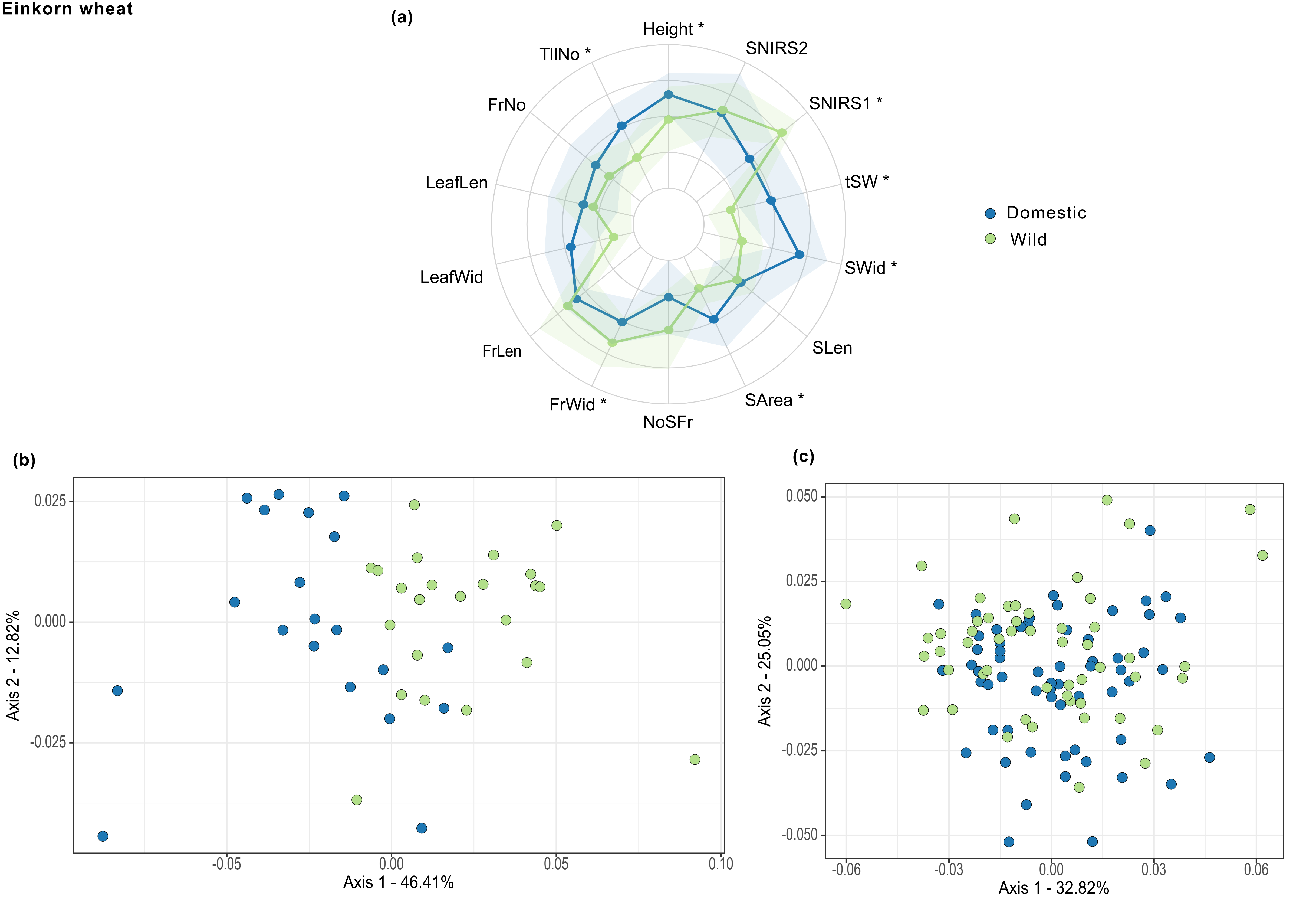

### Figure S7

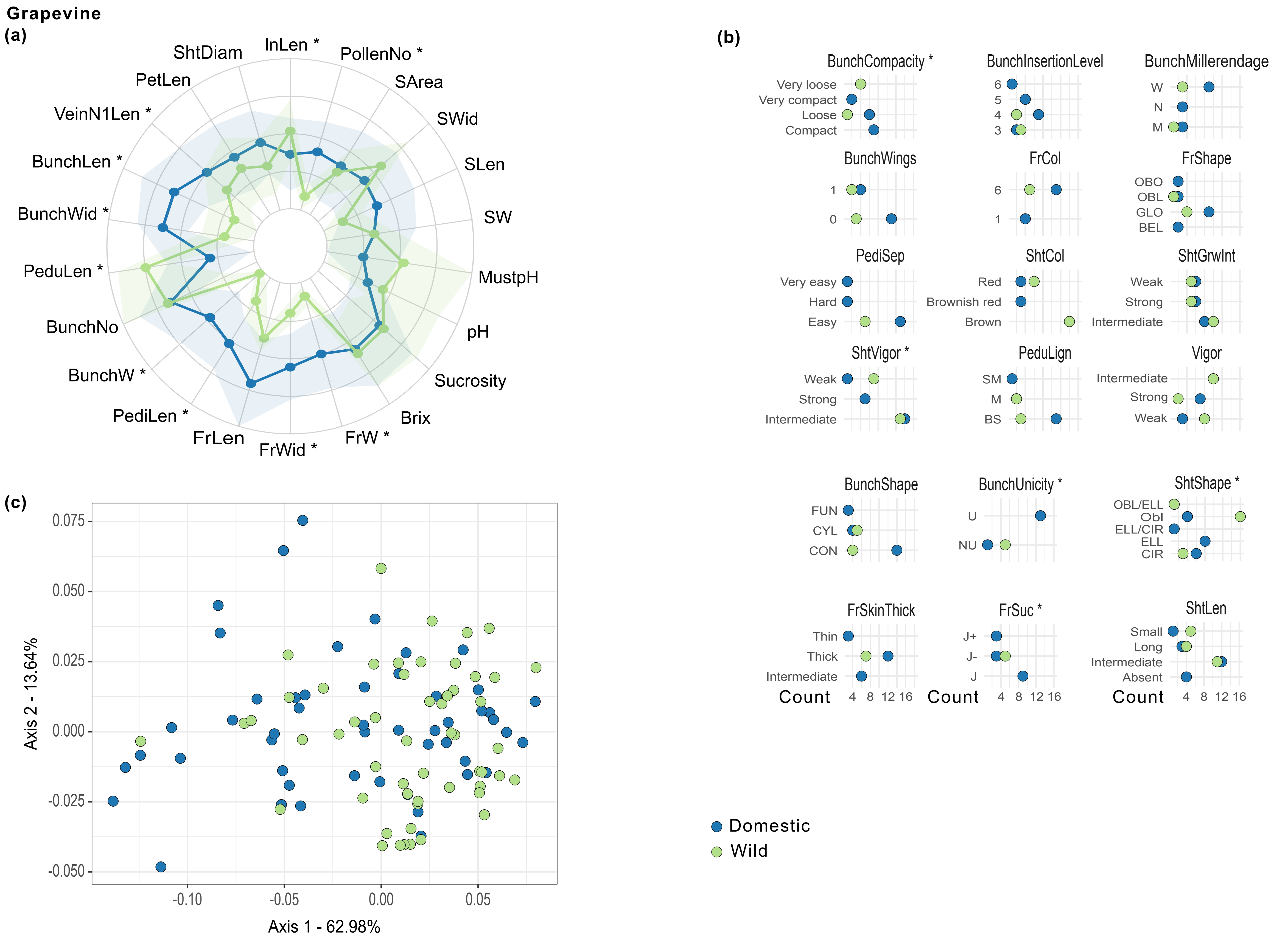

### Figure S8

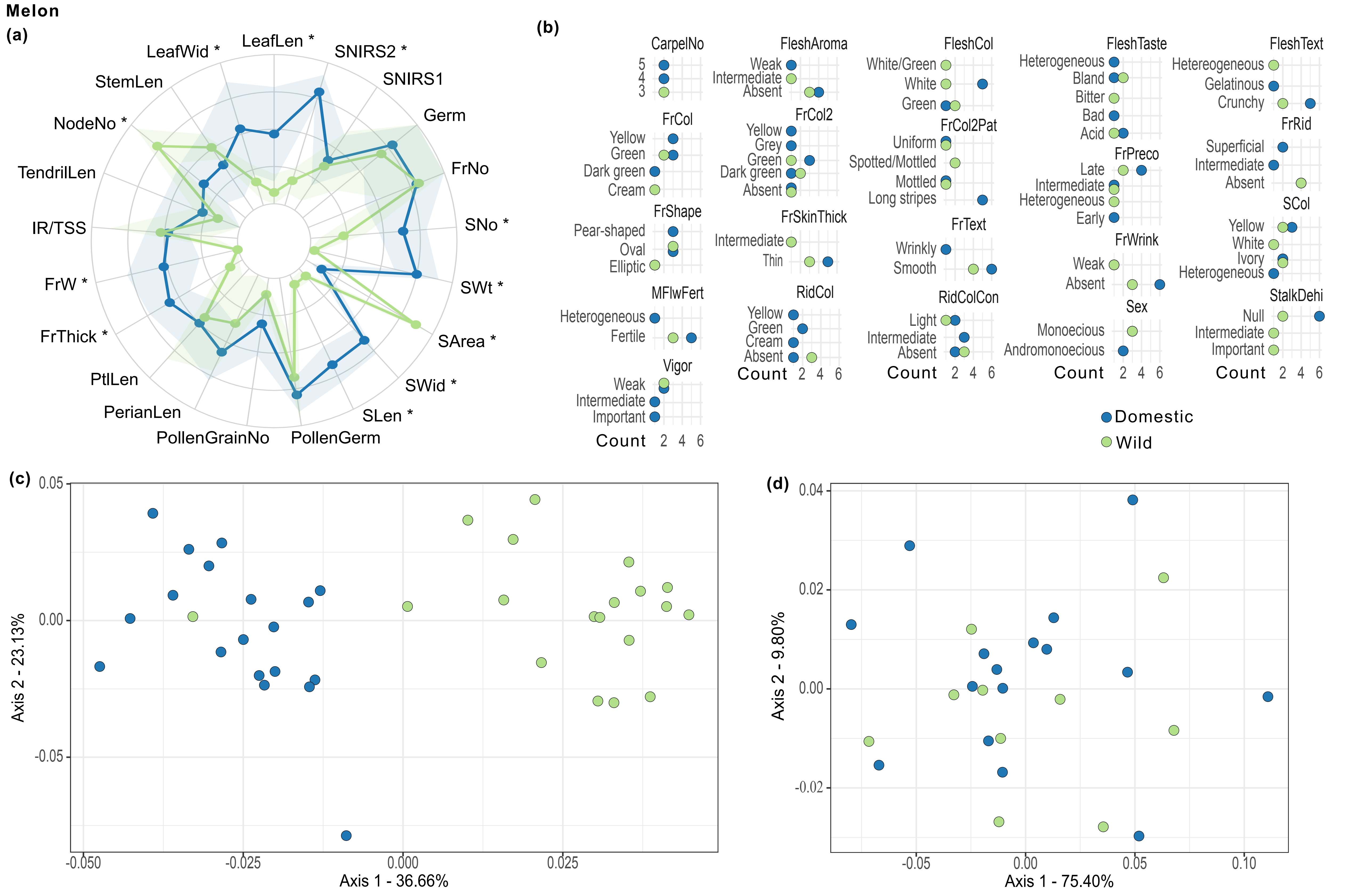

### Figure S9

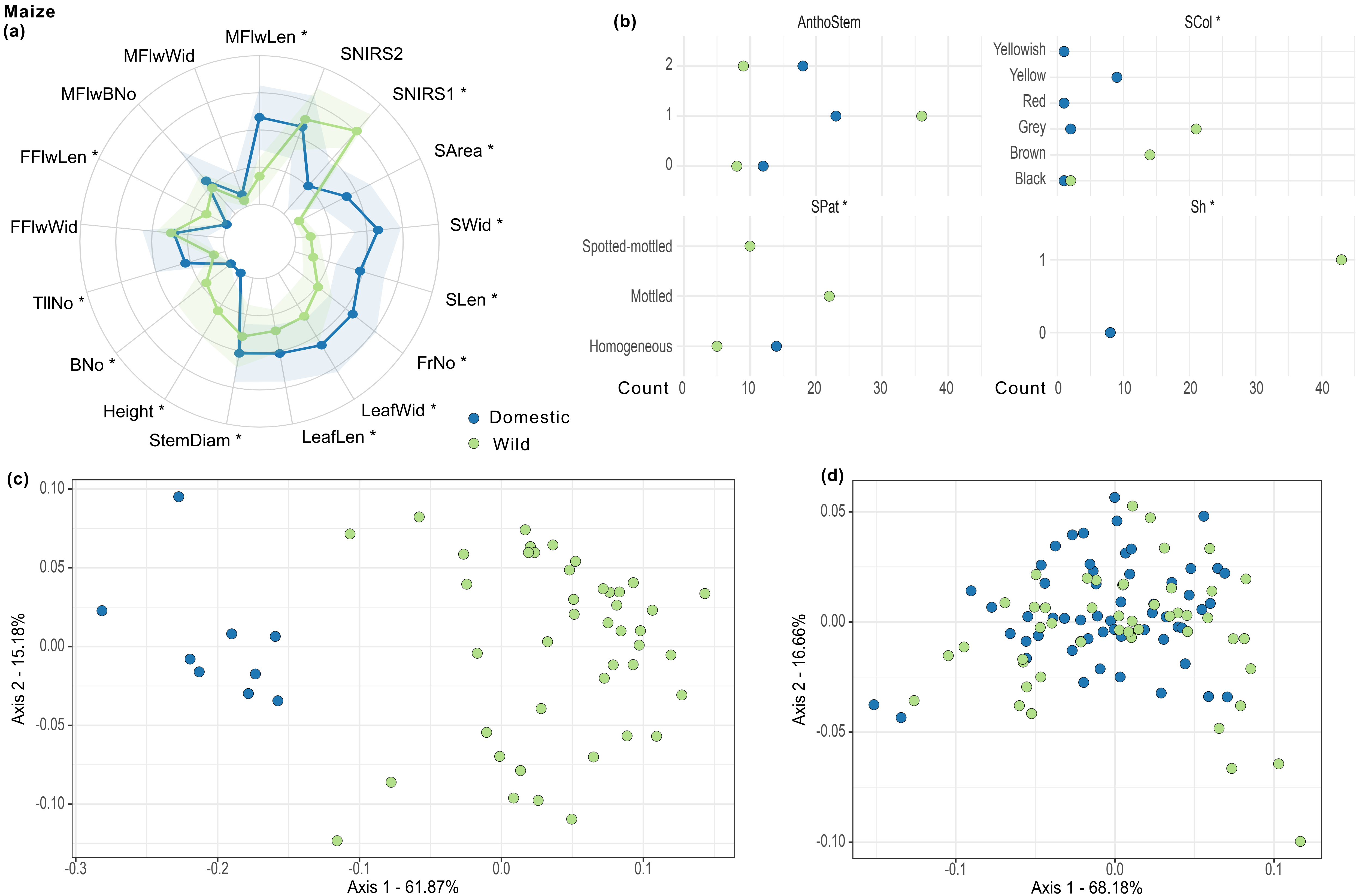

### Figure S10

Pearl millet

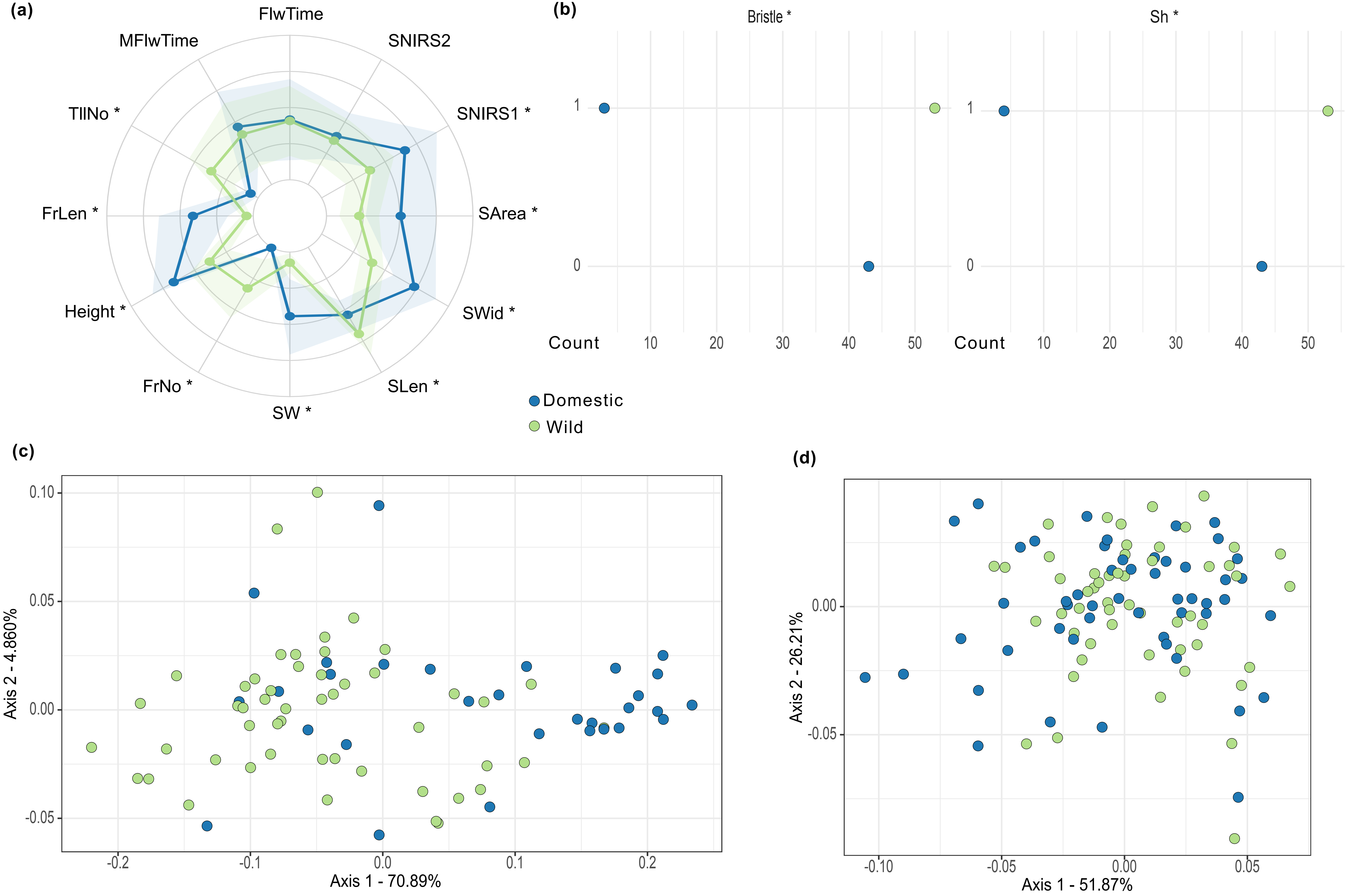

### Figure S11

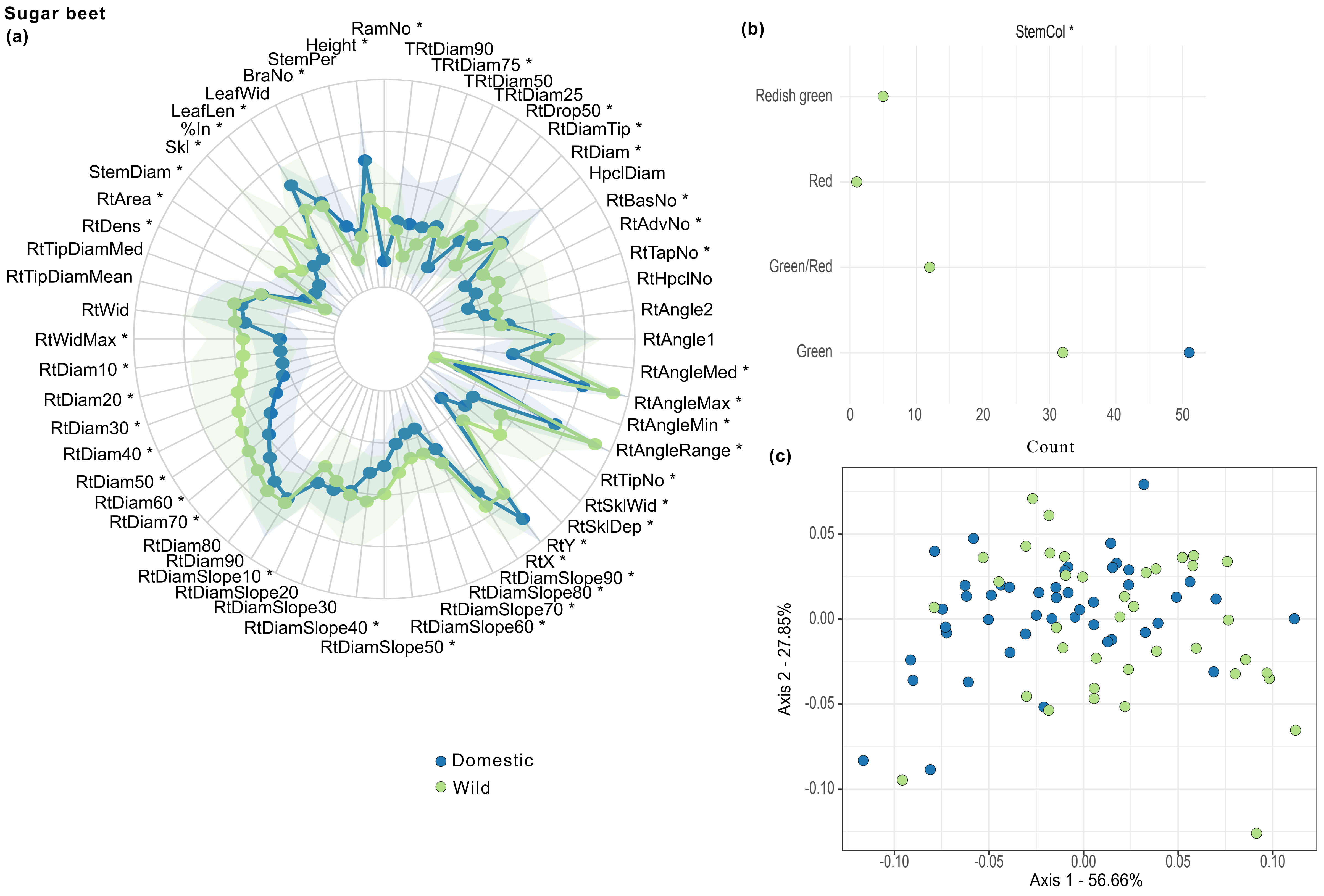

### Figure S12

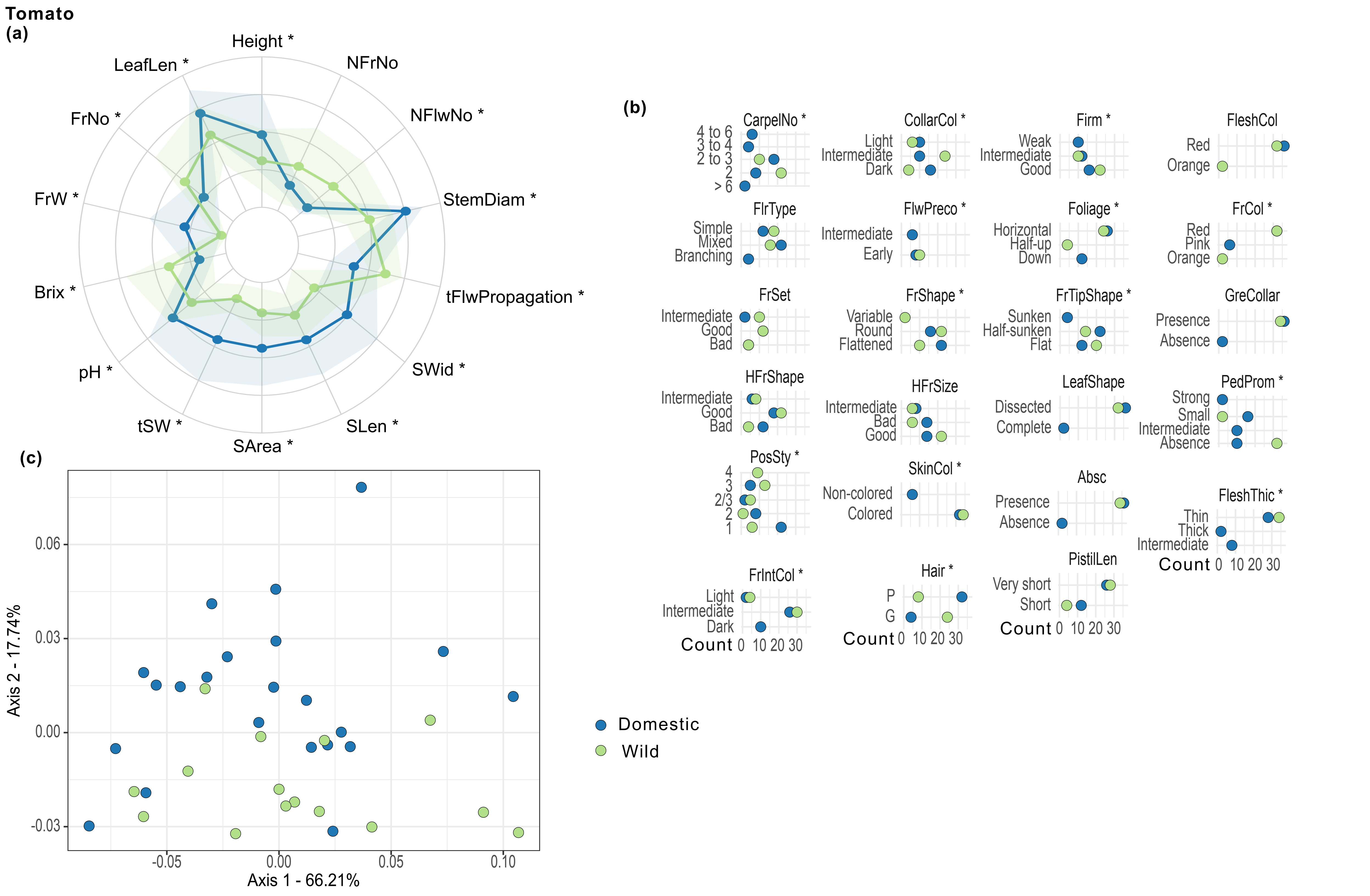

### Figure S13

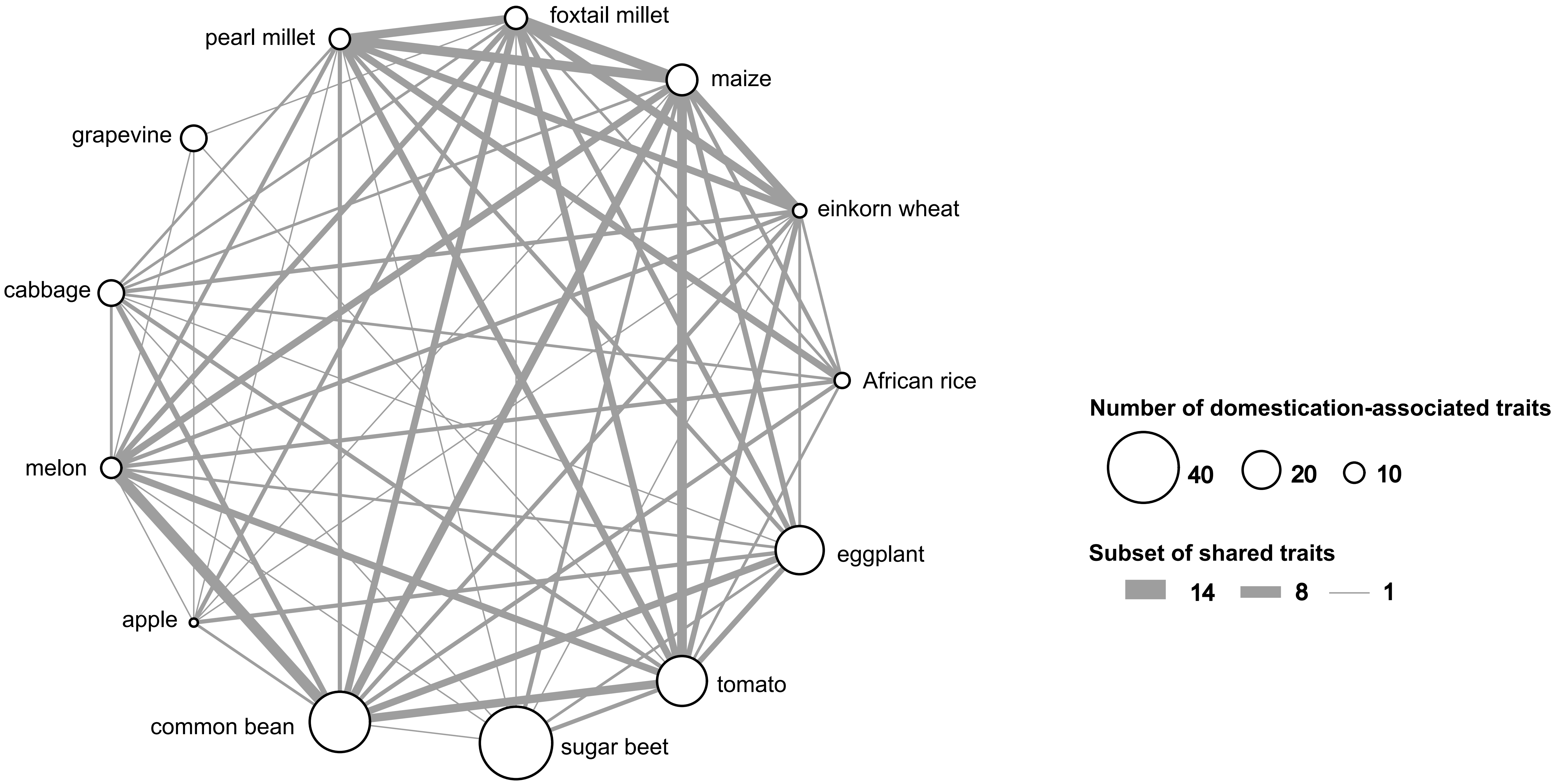

### Figure S14

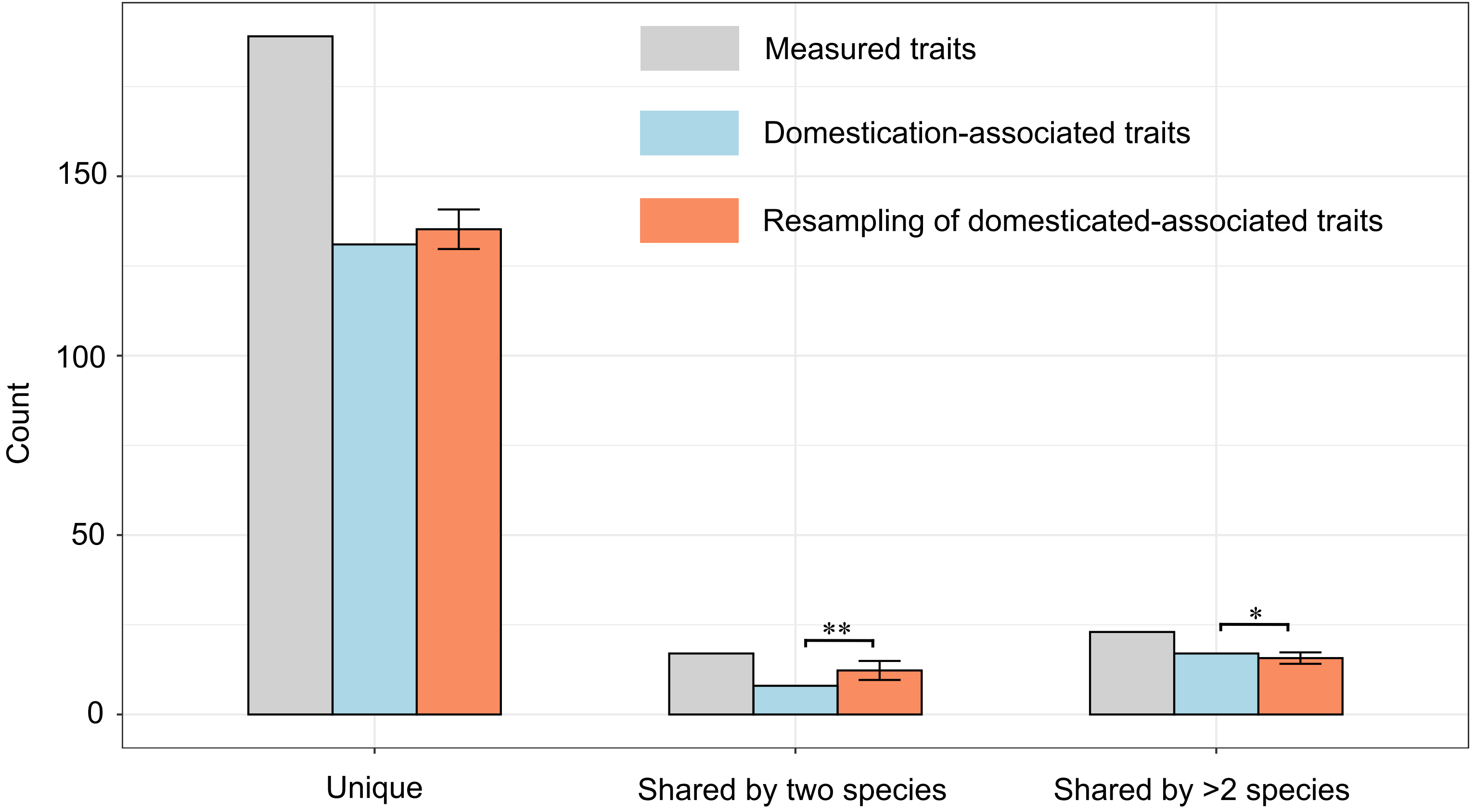

### Figure S15

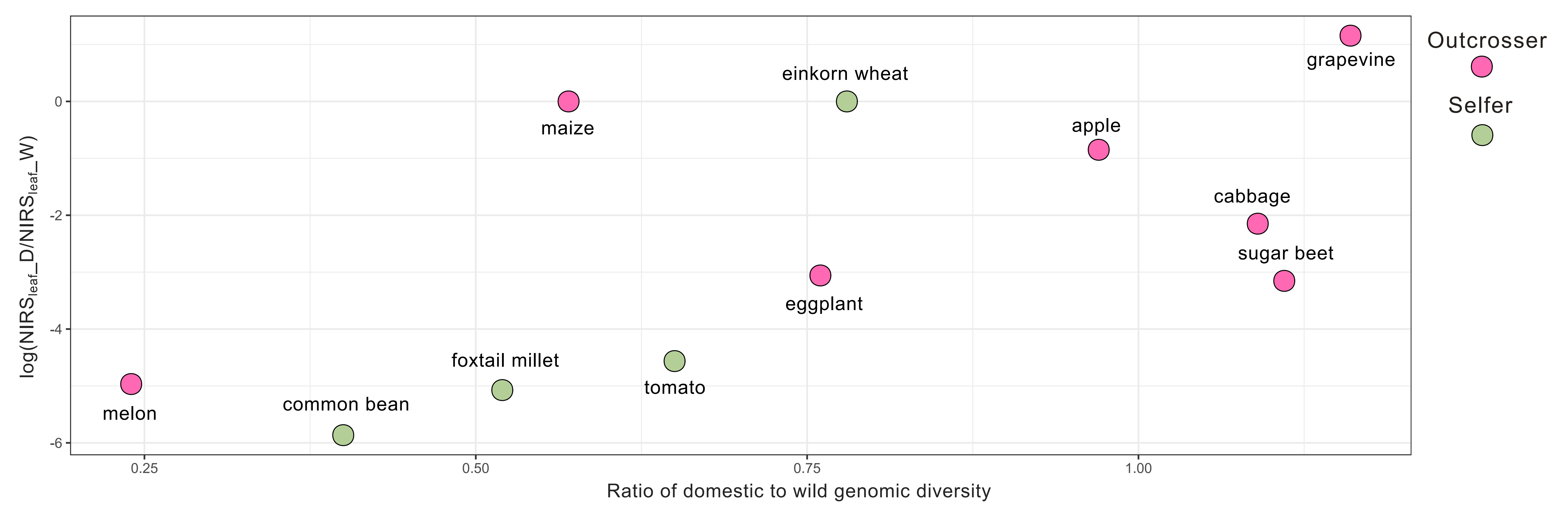
